# The SET1/COMPASS subunit RBBP5 orchestrates epigenetic control of global proteostasis and the 12h oscillator to safeguard metabolic and cellular homeostasis

**DOI:** 10.1101/2024.09.13.612812

**Authors:** Syeda Kubra, Michelle Sun, William Dion, Ahmet Catak, Hannah Luong, Haokun Wang, Yinghong Pan, Jia-Jun Liu, Aishwarya Ponna, Lijun Liu, Ian Sipula, Jian-Hua Luo, Michael J. Jurczak, Silvia Liu, Bokai Zhu

## Abstract

Proteostasis is vital for cellular health, with disruptions leading to aging, neurodegeneration and metabolic disorders. Traditionally, proteotoxic stress responses were studied as acute reactions to various noxious factors; however, recent evidence reveals that many proteostasis genes exhibit ∼12h ultradian rhythms under physiological conditions in mammals, driven by an XBP1s-dependent 12h oscillator. By examining the chromatin landscape of this oscillator, we identified RBBP5 as an essential epigenetic regulator of global proteostasis dynamics. Mechanistically, as the core subunit of the SET1/COMPASS complex, RBBP5 co-activates XBP1s to facilitate dynamic proteostasis gene expression by marking promoter-proximal H3K4me3, which further recruits the Integrator Complex and SWI/SNF chromatin remodelers. Functionally, RBBP5 is indispensable for regulating both the 12h oscillator and acute transcriptional response to various proteotoxic stresses, including ER stress and nutrient deprivation. RBBP5 ablation causes increased susceptibility to proteotoxic stress, chronic inflammation, and hepatic steatosis in mice, along with impaired autophagy and reduced cell survival *in vitro*. In humans, lower *RBBP5* expression is associated with reduced adaptive stress-response gene expression and hepatic steatosis. Our findings not only highlight a previously unrecognized epigenetic timing mechanism distinct from circadian regulation but also establish RBBP5 as a central regulator of proteostasis, essential for cellular resilience and organismal health.

**One sentence summary:** RBBP5 regulates global mammalian proteostasis.

## Introduction

Proteostasis, the maintenance of proper protein folding, trafficking, and turnover, is a major challenge for the cell (1). Much of the cellular energy expenditure is dedicated to maintaining a healthy proteome, as millions of molecules are produced every single minute (2). Challenges to proteostasis, exemplified by the accumulation of unfolded or misfolded proteins, can lead to a range of cellular dysfunctions and pathological conditions, such as aging, neurodegenerative diseases, cancer, and metabolic disorders (1,3–6). To cope with proteotoxic stress, organisms have developed a wide range of stress response mechanisms, including the heat shock response, the unfolded protein response (UPR), and the more generalized integrated stress response (ISR) (7–9). Although the detailed signaling molecules vary, these pathways generally involve transmitting the information of cellular stress to the nucleus by activating various transcription factors (TF) containing the basic leucine zipper domain (bZIP), such as spliced form of XBP1 (XBP1s), ATF4 and ATF6 and heat shock factor 1 (HSF1). After binding the stress response element in the promoters/enhancers, these TFs in turn upregulate a plethora of genes aimed to restore proteostasis (10,11).

While proteotoxic stress responses were traditionally viewed and studied as distinct acute responses to different “noxious” factors, recent evidence by our group indicates that at the physiological level (and in the absence of evident extrinsic stress), the expression of hundreds of proteotoxic stress response regulatory and output genes exhibit cell-autonomous ultradian rhythms cycling with a ∼12h period in multiple mammalian cell-types and tissues, including humans (12–21). These proteostasis genes encompass both output genes involved in protein processing in the ER/Golgi, redox regulation, protein folding, ER-associated protein degradation (ERAD), autophagy, and upstream regulatory molecules and TFs like *Xbp1*, *Atf4*, *Atf6*, *Ddit3*, *Perk* and *Ire1α* (12,22). Many of these ∼12h ultradian rhythms are established by an XBP1s-dependent 12h ultradian oscillator separate from both the ∼24h circadian clock and the cell cycle (21,23,24). It is hypothesized that the mammalian 12h oscillator may have evolved from the circatidal clock of marine animals, and later co-opted to adapt to the ∼12h cycle of metabolic stress peaking at transition times at dawn and dusk in terrestrial animals (13,16,21,25). The 12h oscillator is well-studied in the liver, and mice with liver-specific 12h oscillator ablation exhibited accelerated liver aging and steatosis (19), manifested with impaired proteostasis, lipid metabolism, and mitochondria dysfunction (19,20,26). Since the 12h oscillator integrates multiple proteotoxic stress signaling, it provides us with a unique opportunity to study proteostasis using a holistic approach: rather than studying individual stress response separately, by investigating the 12h oscillator we could gain new mechanistic insights for universal proteostasis regulation.

While significant work has been done on upstream proteotoxic stress sensing and protein-folding mechanisms in the ER and cytosol, there remains a lack of fundamental understanding of the transcriptional regulation of proteostasis in general, particularly concerning the temporal epigenome dynamics, chromatin landscapes, and co-regulatory networks underlying global proteostasis control. In this study, we seek to address an important unanswered question: is there a designated epigenetic regulator that orchestrates global proteostasis dynamics? By analyzing the chromatin landscape of the murine 12h hepatic oscillator, we identified RBBP5—an essential subunit of the Complex Proteins Associated with Set1 (COMPASS) complex responsible for depositing H3K4me3 (27) —as a pivotal epigenetic regulator that governs global proteostasis dynamics. By contrast, histone acetyltransferase (HAT) and associated H3K9Ac are dispensable for dynamic proteostasis gene expression. RBBP5 plays a critical role in regulating both the 12h oscillator and transcriptional response to a plethora of acute proteostasis stress, functioning as a co-activator for XBP1s, to mark H3K4me3 at the promoter of proteostasis genes. Proximity labeling of H3K4me3 further reveals the chromatin-associated proteomic architecture during proteostatic stress, including subunits of the Integrator complex and SWI/SNF chromatin remodelers. Loss of RBBP5 leads to heightened susceptibility to proteotoxic stress, driving chronic inflammation and hepatic steatosis in mice, while also impairing autophagy and cell survival *in vitro*. In humans, reduced RBBP5 expression is linked to a weakened adaptive stress response and the onset of hepatic steatosis. Our findings not only highlight a previously unrecognized epigenetic timing mechanism distinct from circadian regulation but also establish RBBP5 as a central regulator of proteostasis, essential for cellular resilience and organismal health.

## Materials and Methods

### Plasmids

pCDH-EF1a-mCherry-EGFP-LC3B was a gift from Sang-Hun Lee (Addgene plasmid # 170446; http://n2t.net/addgene:170446; RRID: Addgene_170446) (28). pLenti-X1-hygro-mCherry-RAMP4 was a gift from Jacob Corn (Addgene plasmid # 118391; http://n2t.net/addgene:118391; RRID:Addgene_118391) (29).

### The generation of *Rbbp5* ^(flox/flox)^ mice

The Rbbp5-flox allele was generated by the Mouse Embryo Services Core (University of Pittsburgh, Department of Immunology). The targeting strategy is based on the Easi-CRISPR method (30). In brief, fertilized embryos from C57BL/6J background, produced by natural mating, were microinjected in the pronuclei with a mixture of 0.67 µM EnGen Cas9 protein (New England Biolabs, Cat.No. M0646T), two Cas9 guides RNA: Rbbp5-695rev and Rbbp5-951forw (21.23 ng/µl each) and a long single stranded oligonucleotides Rbbp5-flox-ssODN (5 ng/µl). The long single stranded oligonucleotide encoding the donor sequence was synthesized by Integrated DNA Technologies (IDT).

To produce the sgRNA, a double strand linear DNA template is created by annealing of the following target specific oligonucleotides with a common primer (gRNA-Scaffold-R: 5’-AAAAAAGCACCGACTCGGTGCCACTTTTTCAAGTTGATAACGGACTAGCCTTATTTTAACTTG CTATTTCTAGCTCTAAAAC-3’) containing the full tracrRNA sequence and then PCR amplified as

previously described (31). The sgRNA was synthesized using HiScribe T7 Quick High Yield RNA Synthesis Kit (New England Biolabs, Cat. No. E2050S).

T7-Rbbp5-695rev sequence:

TAATACGACTCACTATAGGAAGCATTCTAAGATTAAACCGTTTTAGAGCTAGAAATAGCA

T7-Rbbp5-951forw sequence:

TAATACGACTCACTATAGGAGTAACAGATAGTATTCCGAGTTTTAGAGCTAGAAATAGCA

Rbbp5-flox-ssODN sequence:

gcagttaataaacttgaagtgtttatatagaaaatacatcaggatggcaaccaaaccctttagatacctgagattttattctcttatcccaggtATAACTTCGTATAGCATACATTATACGAAGTTATgaattcttaatcttagaatgcttagtctgtttcatgctgaagataagttgtcagaactcacattgttactgatttttcgttcagtgactcgtctgttttgtcataggaaaattggagtgcatttgcaccagacttcaaagagttggatgaaaatgtagaatatgaggaaagagaatcagaatttgatattgaggatgaagataagagtgagcctgagcaaacaggtgatgcctctcagtaacagatagtattgaattcATAACTTCGTATAGCATACATTATACGAAGTTATccgaagggaaggactgtcttactcaatgctctttgaattagtgaatgcatttatttactttgtgttgtaactgtgctcaaagcttcatctgccacgtttccctcacttgtacatagttttcacagtgcttgaagagtcttactg

The injected zygotes were cultured overnight, the next day the embryos that developed to the 2-cell stage were transferred to the oviducts of pseudo pregnant CD1 female surrogates. Following optimization of the cycling conditions, the potential founder mice were genotype with the following primers pair, using Q5 High-Fidelity DNA Polymerase (NEW ENGLAND BIOLABS, Cat. no. M0491L). The body weight of male adult mice was measured weekly.

*Rbbp5* genotyping forward primer: AGTTCAGAGCTGGTTTCCAAC

*Rbbp5* genotyping reverse primer: AGGTAATGCCCATGTGAGCC

### Biological rhythm study in mice

All mice used for biological rhythm study are in C57BL/6J background, male and between 3 and 4 months of age. Wild-type C57BL/6J mice (n=12), *Rbbp5* (flox/flox); Alb-CRE (-/-) (n=48) and *Rbbp5* (flox/flox); Alb-CRE (+/-) mice (n=48) [*Rbbp5* (flox/flox); Alb-CRE (-/-) and *Rbbp5* (flox/flox); Alb-CRE (+/-) mice were littermates] were first entrained under LD12:12 conditions for 2 weeks before transferred to constant darkness for 24hrs. Mice were then sacrificed via cervical dislocation at a 2h interval [for *Rbbp5* (flox/flox); Alb-CRE (-/-) and *Rbbp5* (flox/flox); Alb-CRE (+/-) mice] or 4h interval (for wild-type C57BL/6J mice) for a total of 48 hours under constant darkness. Mice were fed *ad libitum* during the entire experiment. The animal studies were carried out in accordance with the National Institutes of Health guidelines and were granted formal approval by the University of Pittsburgh’s Institutional Animal Care and Use Committee (approval number IS00013119 and IS00013119).

### Food intake and locomotor activity monitoring

Promethion Multi-plexed Metabolic Cage System was used for real-time measuring of food intake and locomotor activity. Male *Rbbp5 ^Flox^* and *Rbbp5 ^LKO^* mice between 3 and 4 months of age were acclimated to the chambers and ad libitum food intake was monitored for 48 hours under LD12:12 followed by 48 hours of constant darkness. n=6 for *Rbbp5 ^Flox^* and n=8 for *Rbbp5 ^LKO^* mice for LD12:12 study. n=5 for *Rbbp5 ^Flox^* and n=3 for *Rbbp5 ^LKO^* mice for DD study.

### ER stress induction study in mice

For acute ER stress induction experiment, male littermates *Rbbp5* (flox/flox); Alb-CRE (-/-) and *Rbbp5* (flox/flox); Alb-CRE (+/-) mice between 5 and 7 months of age were randomly divided into two groups, and intraperitoneally injected with 0.05mg/kg body weight of tunicamycin dissolved in 500ul 3% DMSO in PBS or vehicle control (3% DMSO in PBS), respectively. Mice were injected with Tu between 9∼10am in the morning and 8 hours later dissected for transcriptomic and western blot analysis for different tissues. The sample size is n=3 for *Rbbp5 ^Flox^*DMSO, n=3 for *Rbbp5 ^Flox^* Tu, n=4 for *Rbbp5 ^LKO^*DMSO, n=4 for *Rbbp5 ^LKO^* Tu.

For experiments involving RBBP5 liver-overexpressing mice, AAV8-TBG-EGFP and AAV8-TGB-mRBBP5 viruses were customarily designed and produced from Vector Biolab (Malvern, PA). Male wildtype C57BL/6J mice between 2 and 3 months of age were tail-vein injected with 0.5x10^11 genome copies of AAV8-TBG-EGFP or AAV8-TGB-mRBBP5 virus diluted in 100 µl of PBS. 30 days later, mice injected with either viruses were randomly divided into two cohorts and intraperitoneally injected with 0.05mg/kg body weight of tunicamycin dissolved in 500ul 3% DMSO in PBS or vehicle control (3% DMSO in PBS). Mice were injected with Tu between 9∼10am in the morning and 8 hours later dissected for transcriptomic and western blot analysis for different tissues. The sample size is n=3 for AAV8-TBG-EGFP DMSO, n=4 for AAV8-TBG-EGFP Tu, n=4 for AAV8-TGB-mRBBP5 DMSO and n=4 for AAV8-TGB-mRBBP5 Tu.

For chronic ER stress induction experiment, male littermates *Rbbp5* (flox/flox); Alb-CRE (-/-) and *Rbbp5* (flox/flox); Alb-CRE (+/-) mice between 5 and 7 months of age were intraperitoneally injected with 0.025mg/kg body weight of tunicamycin dissolved in 500ul 3% DMSO in PBS or vehicle control (3% DMSO in PBS) daily for six consecutive days. Mice were injected with Tu between 9∼10am in the morning and 24 hours after the last injection dissected for transcriptomic, western blot analysis, histology for liver tissues. The sample size is n=4 for *Rbbp5 ^Flox^* DMSO, n=4 for *Rbbp5 ^Flox^* Tu, n=2 for *Rbbp5 ^LKO^* DMSO, n=3 for *Rbbp5 ^LKO^* Tu.

### Oil Red O Staining

Frozen sections were rinsed with PBS and then fixed with 10% neutral buffered formalin for 30 min at RT. After washing twice with deionized water, 60% isopropanol was applied to the fixed cells for 5 min, followed by a freshly prepared working solution of 1.5 mg/ml Oil Red O in isopropanol for 5 min at RT. The stained, fixed tissues were then rinsed with tap water until clear, covered with tap water and viewed on a phase contrast microscope. Size and areas of lipid droplet were quantified by customarily written pipelines in CellProfiler (v4.1.3).

### Generation of stable autophagy reporter and ER-phagy reporter cell line

Primary MEFs were isolated from *Rbbp5* (flox/flox); Rosa26 CRE (+/+) mice and immortalized with SV40 lentivirus as previously described (32). For mCherry-EGFP-LC3B-expressing autophagy reporter MEFs, lentiviruses packaged in HEK293T cells with co-transfection of pCDH-EF1a-mCherry-EGFP-LC3B, pMD2.G and psPAX2 plasmids were used to infect *Rbbp5* (flox/flox); Rosa26 CRE (+/+) MEFs with a multiplicity of infection (MOI) of 3 for three times to achieve near complete infection. For mCherry-RAMP4-expressing ER-phagy reporter MEFs, lentiviruses packaged in HEK293T cells with co-transfection of pLenti-X1-hygro-mCherry-RAMP4, pMD2.G and psPAX2 plasmids were used to infect *Rbbp5* (flox/flox); Rosa26 CRE (+/+) MEFs with a multiplicity of infection (MOI) of 3. Stable MEFs were selected in the presence of 200μg/ml hygromycin.

### siRNA Transient Transfections

Immortalized MEFs isolated from wild-type C57BL/6J mice were transfected with 10µM of different siRNAs for 24∼48 hours with Lipofectamine RNAiMAX reagents (Life technologies) per the manufacturer’s instructions. Source of siRNA are as follows: siGENOME non-targeting siRNA pool (Dharmacon, D-001206-1305), siGENOME SMARTpool mouse *Rbbp5* siRNA (Dharmacon, L-054560-01-0005), siGENOME SMARTpool mouse *Ash2l* siRNA (Dharmacon, L-048754-01-0005), siGENOME SMARTpool mouse *Setd1a* siRNA (Dharmacon, L-051358-01-0005), siGENOME SMARTpool mouse *Kat2a/Gcn5* siRNA (Dharmacon, L-040665-01-0005), siGENOME SMARTpool mouse *Kat2b/Pcaf* siRNA (Dharmacon, L-049885-01-0005) and siGENOME SMARTpool mouse *Xbp1* siRNA (Dharmacon, L-040825-00-0005). MEFs were transfected with 10µM of different siRNAs for 48 hours and treated with DMSO or 100ng/ml of tunicamycin for 8h.

### Real-time Luminescence Assay

Stable *Manf*-dluc MEFs were transfected with non-targeting, *Rbbp5,* or *Ash2l* siRNA for 48 hours in DMEM (4.5g/L glucose) supplemented with 10% FBS and treated with 25ng/ml tunicamycin in DMEM for 2h before subjected to real-time luminescence assay using a Lumicycle (Actimetrics) as previously described (33). Briefly, after tunicamycin treatment, MEFs were washed with 1x PBS and cultured with DMEM (4.5g/L glucose) supplemented with 0.1 mM luciferin and 10mM HEPES buffer in 35 mm tissue culture dishes in the absence of serum and transferred immediately to Lumicycle for real-time luminescence analysis. Periods of oscillation were identified by embedded Periodogram function.

### Immunofluorescence (IF)

Immunofluorescence was performed as previously described (17). Briefly, liver OCT sections were fixed in cold acetone for 10 mins at -20 °C. The sections were then air dried, rehydrated with PBS and permeabilized with PBS+ 0.1% Triton X-100. The sections were then blocked with 10% goat serum at room temperature for 1 hour. Primary antibody against RBBP5 (Bethyl, A300-109A) was added to the OCT section at 1:1000 dilution overnight at 4 °C. Next day, sections were washed three times with PBS and stained with Alexa-555 anti-rabbit secondary antibody at room temperature for 2 hours. After that, the sections were washed with PBS three times and counterstained with DAPI before mounting (with ProlongGold Glass) and imaging using Leica SP8 lightening confocal microscope (Leica Microsystems).

### Time-lapse microscopy for autophagy reporter MEFs

Time-lapse imaging was performed using SP8 lightening confocal microscope (Leica Microsystems) with Okolab stage top incubator to maintain constant CO_2_ (5%), temperature (37 °C) and humidity (90%). mCherry-EGFP-LC3B-expressing *Rbbp5* (flox/flox); Rosa26 CRE (+/+) MEFs were plated into 8-well chamber slides in full DMEM media and treated with (to deplete RBBP5) or without tamoxifen (3µM/ml) for seven days before replacing with full DMEM or leucine-free media counterstained with DAPI and images were taken every 2 hours for a total of 16 hours. The number of cytosolic autophagosomes (GFP and mCherry double positive foci) and autolysosomes (GFP negative and mCherry positive foci) per cell were quantified manually.

### Immunoblot

Nuclear extracts were made from liver according to previously published protocol (34). Whole cell lysates were lysed in RIPA buffer as previously described (17). Protein concentrations were determined by Bradford assays (Bio-Rad), and aliquots were snap-frozen in liquid nitrogen and stored at -80°C until usage. Immunoblot analyses were performed as described previously (35). Briefly, 25∼50µg proteins separated by 4∼20% gradient SDS-PAGE gels (Biorad) were transferred to nitrocellulose membranes, blocked in TBST buffer supplemented with 5% bovine serum albumin (BSA) or 5% fat-free milk and incubated overnight with primary antibody anti-PERK (Cell signaling, #3192), anti-phospho-PERK (Thr908) (Thermo Fisher, MA5-15033), anti-ATF4 (Cell signaling, #11815), anti-IRE1α (Cell signaling, #3294), anti-phospho-IRE1α (Ser724) (ABclonal, AP0878), anti-XBP1s (Biolegend, 658802), anti-RBBP5 (Bethyl, A300-109A), anti-RBBP5 (D3I6P) (Cell signaling, #13171), anti-WDR5 (Cell signaling, #13105), anti-ASH2L (Bethyl, A300-489A), anti-ASH2L (Bethyl, A300-112A), anti-SETD1A (Diagenode, CS-117-100), anti-SETD1B (Diagenode, CS-118-100), anti-MLL1-c (D6G8N) (Cell signaling, #14197), anti-mCherry (E5D8F) (Cell signaling, #43590), anti-p62/SQSTM1 (Santa Cruz, sc-28359) and anti-β-ACTIN (Cell signaling, #12620) at 4°C. Blots were incubated with an appropriate secondary antibody coupled to horseradish peroxidase at room temperature for 1 hour, and reacted with ECL reagents per the manufacturer’s (Thermo) suggestion and detected by Biorad ChemiDoc MP Imaging System.

### Co-Immunoprecipitation (Co-IP)

Nuclear extracts (NE) were made from liver according to our published protocol (34). Protein concentrations were determined by Bradford assays (Bio-Rad), and aliquots were snap-frozen in liquid nitrogen and stored at -80°C until usage. 200 μg of NE was used for per IP. 5 μg of different antibodies or control IgG were coupled to 1.5 mg of magnetic Dynabeads (Life Technologies) for every IP using Dynabeads Antibody Coupling Kit (Life Technologies) per manufactures’ protocol. Co-IP was essentially carried out as previously described (36) except that coupled antibody Dynabeads was added to the NEs for incubation. The antibodies used for Co-IP are as follows: anti-XBP1s (Biolegend, 658802).

### qRT-PCR

Total mRNA was isolated from murine embryonic fibroblasts (MEFs) or mouse liver with PureLink RNA mini kit (Life Technologies) with additional on-column DNase digestion step to remove genomic DNA per the manufacturer’s instructions. Reverse transcription was carried out using 5µg of RNA using Superscript III (Life Technologies) per the manufacturer’s instructions. For gene expression analyses, cDNA samples were diluted 1/30-fold (for all other genes except for 18sRNA) and 1/900-fold (for 18sRNA). qPCR was performed using the SYBR green system with sequence-specific primers. All data were analyzed with 18S or *β-actin* as the endogenous control. qPCR primer sequences are as follows and all primers span introns, except for primers for quantifying pre-mRNAs:

Mouse *Rbbp5* forward primer: CAGAGCCTATGCAGAAGCTGCA

Mouse *Rbbp5* reverse primer: CTTCACCAGGTTGCCAATGCTC

Mouse *Ddit3* forward primer: GGAGGTCCTGTCCTCAGATGAA

Mouse *Ddit3* reverse primer: GCTCCTCTGTCAGCCAAGCTAG

Mouse *Creld2* forward primer: CTACACCAAGGAGAGTGGACAG

Mouse *Creld2* reverse primer: TCTGTCTCCTCAAAGCCTTCCG

Mouse *Hspa5* forward primer: TGTCTTCTCAGCATCAAGCAAGG

Mouse *Hspa5* reverse primer: CCAACACTTCCTGGACAGGCTT

Mouse *Dnajb11* forward primer: TGTGACCGTCTCACTGGTTGAG

Mouse *Dnajb11* reverse primer: CCCTTTCTTCCACAGCTTGGCT

Mouse *Pdia4* forward primer: TGGGCTCTTTCAGGGAGATGGT

Mouse *Pdia4* reverse primer: GGGAGACTTTCAGGAACTTGGC

Mouse *Hsp90b1* forward primer: GTTTCCCGTGAGACTCTTCAGC

Mouse *Hsp90b1* reverse primer: ATTCGTGCCGAACTCCTTCCAG

Mouse *Syvn1* forward primer: CCAACATCTCCTGGCTCTTCCA

Mouse *Syvn1* reverse primer: CAGGATGCTGTGATAAGCGTGG

Mouse *Gdf15* forward primer: ACTCAGTCCAGAGGTGAGATTG

Mouse *Gdf15* reverse primer: GGGGCCTAGTGATGTCCCAG

Mouse *Il1a* forward primer: ACGGCTGAGTTTCAGTGAGACC

Mouse *Il1a* reverse primer: CACTCTGGTAGGTGTAAGGTGC

Mouse *Il1b* forward primer: TGGACCTTCCAGGATGAGGACA

Mouse *Il1b* reverse primer: GTTCATCTCGGAGCCTGTAGTG

Mouse *Fga* forward primer: GGATTCTAACTCACTGACCAGGA

Mouse *Fga* reverse primer: CCTCAGGATCTCAATTCTGCGC

Mouse *C3* forward primer: CGCAACGAACAGGTGGAGATCA

Mouse *C3* reverse primer: CTGGAAGTAGCGATTCTTGGCG

Mouse *Tifa* forward primer: AGCAAGAAAACCAGTTTGATGGTAG

Mouse *Tifa* reverse primer: GCAACAGGAACTGATACTCTCCG

Mouse *Fgl1* forward primer: GGAAACTGTGCTGAGGAAGAGC

Mouse *Fgl1* reverse primer: TCCGTTTCTGCCCTGTAGGAAC

Mouse *Fgb* forward primer: TGACACCTCCATCAAGCCGTAC

Mouse *Fgb* reverse primer: GGTCCCATTTCCTGCCAAAGTC

Mouse *Il6* forward primer: CTTCCATCCAGTTGCCTTCT

Mouse *Il6* reverse primer: CTCCGACTTGTGAAGTGGTATAG

Mouse *Herpud1* forward primer: acctgagccgagtctaccc

Mouse *Herpud1* reverse primer: aacagcagcttcccagaataaa

Mouse *Nbr1* forward primer: GGATTTAAAGCACCTCCTGATTCC

Mouse *Nbr1* reverse primer: ATTGGGTCCCACTCAGGTCT

Mouse *Sesn2* forward primer: CTGGCCGAACTCATCCAAG

Mouse *Sesn2* reverse primer: CTGCCTCATGCGTTCCATC

Mouse *Sqstm1* forward primer: AACAGATGGAGTCGGGAAAC

Mouse *Sqstm1* reverse primer: AGACTGGAGTTCACCTGTAGA

Mouse *Stbd1* forward primer: CTGGGAAGTAGATGGGAAAGTG

Mouse *Stbd1* reverse primer: TCGGTGTTTGTGGTGTAGTG

Mouse *Sirt1* forward primer: TGACAGAACGTCACACGCCA

Mouse *Sirt1* reverse primer: ATTGTTCGAGGATCGGTGCCA

Mouse *Gcn5* forward primer: TTGATTGAGCGCAAACAGGC

Mouse *Gcn5* reverse primer: CAGCCTGTCTCTCGAATGCC

Mouse *Pcaf* forward primer: CGTGAAGAAGGCGCAGTTG

Mouse *Pcaf* reverse primer: CAGGACTCCTCTGCCTTGC

Mouse *Ash2l* forward primer: gctgtgtctgctagtgggaac

Mouse *Ash2l* reverse primer: catcttgctgcttacgcttg

Mouse *Setd1a* forward primer: gggtagcaccccctactctc

Mouse *Setd1a* reverse primer: gggtttgaaggaggttgaagt

Mouse *Eif2ak3* forward primer: ccttggtttcatctagcctca

Mouse *Eif3ak3* reverse primer: atccagggaggggatgat

Mouse total *Xbp1* forward primer: gggtctgctgagtcc

Mouse total *Xbp1* reverse primer: cagactcagaatctgaagagg

Mouse *Sec23b* forward primer: tgaccaaactggacttctgga

Mouse *Sec23b* reverse primer: aaagaatctcccatcaccatgt

Mouse *Manf* forward primer: gacagccagatctgtgaactaaaa

Mouse *Manf* reverse primer: tttcacccggagcttcttc

Mouse *Hyou1* forward primer: GAGGCGAAACCCATTTTAGA

Mouse *Hyou1* reverse primer: GCTCTTCCTGTTCAGGTCCA

Mouse *Atf4* forward primer: CCACTCCAGAGCATTCCTTTAG

Mouse *Atf4* reverse primer: CTCCTTTACACATGGAGGGATTAG

Mouse *Atf6* forward primer: CATGAAGTGGAAAGGACCAAATC

Mouse *Atf6* reverse primer: CAGCCATCAGCTGAGAATTAGA

Mouse 18s RNA forward primer: ctcaacacgggaaacctcac

Mouse 18s RNA reverse primer: cgctccaccaactaagaacg

Mouse *β-actin* forward primer: aaggccaaccgtgaaaagat

Mouse *β-actin* reverse primer: gtggtacgaccagaggcatac

### RNA-Seq to identify RBBP5-dependent 12h hepatic transcriptome

Mouse liver tissues were collected from *Rbbp5 ^Flox^* (n=2) and *Rbbp5 ^LKO^* (n=2) mice at 2h intervals for a total of 48 hours under constant darkness condition. Total RNA was isolated from mouse liver with miRNeasy Mini Kit (Qiagen) per the manufacturer’s instructions. RNA was assessed for quality using an Agilent TapeStation 4150/Fragment Analyzer 5300 and RNA concentration was quantified on a Qubit FLEX fluorometer. Libraries were generated with the Illumina Stranded mRNA Library Prep kit (Illumina: 20040534) according to the manufacturer’s instructions. Briefly, 200 ng of input RNA was used for each sample. Following adapter ligation, 12 cycles of indexing PCR were completed, using IDT for Illumina RNA UD Indexes (Illumina: 20040553-6). Library quantification and assessment was done using a Qubit FLEX fluorometer and an Agilent TapeStation 4150/Fragment Analyzer 5300. The prepared libraries were pooled and sequenced using NoveSeq 6000 (Illumina), generating an average of 40 million 2×100 bp paired end reads per sample. RNA-Seq library preparation and sequencing were performed at UPMC Genome Center. Raw RNA-seq FASTQ files were analyzed by FastQC for quality control. Adaptors and low-quality reads were filtered by Trimmomatic (37). Then the processed reads were aligned by HISAT2 (38) against mouse reference mm10. Gene and isoform FKPM values were calculated by Cufflinks. Since the background gene expression level is FPKM=0.1 in mouse liver (21), only those genes with average gene expression across 48 hours in Rbbp5 *^Flox^* mice larger than 0.1 were included for rhythm-identification analysis. Averaged FPKM values at each time were used for cycling transcripts identification by the eigenvalue/pencil method. Raw temporal data was subject to polynomial detrend (n=2) first, and superimposed oscillations were identified using previously described eigenvalue/pencil method on the detrended dataset (23,39). Specifically, three oscillations were identified from each gene. Criterion for circadian gene is period between 21h to 27h, decay rate between 0.9 and 1.1; for ∼12hr genes: period between 10h to 13h, decay rate between 0.9 and 1.1. The relative amplitude is calculated via dividing the amplitude by the mean expression value of each gene. All analysis were performed in MatlabR2017A. Heat maps were generated by Gene Cluster 3.0 and TreeView 3.0 alpha 3.0 using log2 mean-normalized values. RAIN analysis was performed as previously described in Bioconductor (3.4) (http://www.bioconductor.org/packages/release/bioc/html/rain.html) (40) using the full detrended dataset with duplicated at each time point.

For the eigenvalue method, every gene consists of multiple superimposed oscillations. Therefore, we define a circadian gene as any gene that exhibits a superimposed circadian rhythm, regardless of its relative amplitude compared to other superimposed oscillations. Similar definitions apply to 12hr genes. Under this definition, a gene can be both a circadian and 12hr-cycling gene. By comparison, we define a dominant circadian gene as any gene whose superimposed circadian rhythm has the largest amplitude among all oscillations. Similar definitions also apply to 12hr genes. Under this definition, dominant circadian and dominant 12hr genes are mutually exclusive.

### RNA-seq for identifying RBBP5-dependent acute transcriptional response to ER stress

Mouse liver tissues were collected from *Rbbp5 ^Flox^* and *Rbbp5 ^LKO^* mice intraperitoneally treated with or without Tu for 8 hours. Total mRNA was isolated from mouse liver with PureLink RNA mini kit (Life Technologies) with additional on-column DNase digestion step to remove genomic DNA per the manufacturer’s instructions. Detailed procedures for transcriptome sequencing were described previously(41) and were supported by the Genomics and Systems Biology Core of the Pittsburgh Liver Research Center. Briefly, the mRNA samples per mouse were processed into short-read libraries using the Bio-Rad SEQuoia Dual Indexed Primers. Next, the circularization was performed using the Element Biosciences Adept Library Compatibility Kit v1.1 based on the manufacturer’s instructions, followed by the quantification using qPCR with the provided standard.

Finally, the libraries were measured by the Element Biosciences AVITI system to sequence paired end reads with 75 bp each. The raw FASTQ files from RNA-seq were trimmed by Trimmomatic (42) to filter out low-quality reads. The surviving reads were then aligned to the mouse mm10 reference genome by STAR aligner(43). Fragments Per Kilobase of transcript per Million mapped reads (FPKM) values per library were quantified by the tool Cufflinks(44). Log 2 normalized fold induction and adjusted p values between *Rbbp5 ^Flox^* DMSO and *Rbbp5 ^LKO^* DMSO, *Rbbp5 ^Flox^* Tu and *Rbbp5 ^LKO^* Tu, *Rbbp5 ^Flox^* DMSO and *Rbbp5 ^Flox^* Tu, and *Rbbp5 ^LKO^* DMSO and *Rbbp5 ^LKO^* Tu were calculated by using the iDEP (integrated Differential Expression and Pathway analysis) online application (45).

### Chromatin Immunoprecipitation (ChIP)-Seq and ChIP-qPCR

ChIP for RBBP5 was performed using anti-RBBP5 antibody (Bethyl, A300-109A) as previously described (21). Briefly, mouse liver samples were submerged in PBS + 1% formaldehyde, cut into small (∼1 mm3) pieces with a razor blade and incubated at room temperature for 15 minutes. Fixation was stopped by the addition of 0.125 M glycine (final concentration). The tissue pieces were then treated with a TissueTearer and finally spun down and washed twice in PBS. Chromatin was isolated by the addition of lysis buffer, followed by disruption with a Dounce homogenizer. The chromatin was enzymatically digested with MNase. Genomic DNA (Input) was prepared by treating aliquots of chromatin with RNase, Proteinase K and heated for reverse-crosslinking, followed by ethanol precipitation. Pellets were resuspended and the resulting DNA was quantified on a NanoDrop spectrophotometer. An aliquot of chromatin (30 μg) was precleared with protein A agarose beads (Invitrogen). Genomic DNA regions of interest were isolated using 4 μg of antibody. Complexes were washed, eluted from the beads with SDS buffer, and subjected to RNase and proteinase K treatment. Crosslinking was reversed by incubation overnight at 65 °C, and ChIP DNA was purified by phenol-chloroform extraction and ethanol precipitation. The DNA libraries were prepared at the University of Pittsburgh and sequenced at Gene by Gene, Ltd per standard protocols. DNA libraries were prepared with Ovation® Ultralow V2 DNA-Seq library preparation kit (NuGen) using 1ng input DNA. The size selection for libraries was performed using SPRIselect beads (Beckman Coulter) and purity of the libraries were analyzed using the High Sensitivity DNA chip on Bioanalyzer 2100 (Agilent). The prepared libraries pooled and sequenced using Nova-Seq 6000 (Illumina), generating ∼30 million 75 bp single-end reads per sample. ChIP-qPCR for MEFs were essentially performed the same way as previously described with anti-RBBP5 (Bethyl, A300-109A) and anti-H3K4me3 (Active Motif, 61379) antibodies, except that the MEFs were directly fixed with 1% formaldehyde before subject to nuclei isolation and chromatin immunoprecipitation. The primers used for ChIP-qPCR are as follows:

Negative control region 1 forward primer: GCAACAACAACAGCAACAATAAC

Negative control region 1 reverse primer: CATGGCACCTAGAGTTGGATAA

*Manf* promoter forward primer: CCCTTAAATGGGTCAACGTCTC

*Manf* promoter reverse primer: GGCGCTAAACCCAAGGAAA

*Xbp1* promoter forward primer: TCCGTACGGTGGGTTAGAT

*Xbp1* promoter reverse primer: ACCTTCTTCTGTGCCTGTG

*Hyou1* promoter forward primer: CCGTGTGGGTACGTCCT

*Hyou1* promoter reverse primer: GACGGCTGCTCCATCCT

*Sesn2* promoter forward primer: ACAGGAGGCCGGGACTA

*Sesn2* promoter reverse primer: CTGGGCTGAAAGGAGTGTCTAT

### ChIP-Seq analysis

The sequences identified were mapped to the mouse genome (UCSC mm10) using BOWTIE function in Galaxy Deeptool (https://usegalaxy.org/) (46). Only the sequences uniquely mapped with no more than 2 mismatches were kept and used as valid reads. PCR duplicates were also removed. Peak calling was carried out by MACS2 (version 2.1.1.20160309) in Galaxy (options -- mfold 5, 50 --pvalue 1e-4), on each ChIP-seq file against input. To account for the different sequencing depths between samples, the BAM files generated from MACS2 were RPKM normalized to sequencing depth using the bamCoverage function in Galaxy Deeptool and the bigwig files were generated accordingly. The relative intensity of each RBBP5 binding site is further calculated via the computeMatrix function with the RPKM normalized bigwig files and bed files from the peak calling as inputs by calculating the area under the curve.

### Proximity labeling

Proximity labeling was performed using Protein A-TurboID as previously described (47). Briefly, target cells were grown to 90-100% confluency in 15 cm dishes, harvested in PBS using a cell scraper, and pelleted by centrifugation. Nuclei was isolated by resuspending the cell pellet in ice-cold HLB buffer, incubating on ice, and washing through sequential centrifugation steps. Isolated nuclei were incubated with digitonin buffer containing ∼2 µg of IgG or H3K4me3 antibody (Active Motif, 61379) for 20 minutes at room temperature with shaking, followed by incubation with 1.4 µg of Protein A-TurboID for 30 minutes at 4°C. After washing, biotinylation was induced by incubating samples in biotinylation buffer (containing ATP, MgCl₂, and 500 µM biotin) for 10 minutes at 37°C. Nuclei were lysed in RIPA buffer at 4°C overnight, sonicated, and centrifuged to collect the supernatant. Streptavidin affinity purification was performed by incubating lysates with M-270 streptavidin beads for 2 hours at room temperature or overnight at 4°C. Beads were washed extensively with RIPA buffer, and bound proteins were eluted by boiling at 100°C for 15 minutes in 70 µL of 1.5X LDS buffer. Samples were clarified by centrifugation and magnetic separation.

For protein identification, half of each sample was separated ∼1.5 cm on a 10% Bis-Tris Novex mini-gel (Invitrogen) using the MES buffer system. The gel was stained with Coomassie, and each lane was excised into ten equally sized segments. Gel pieces were processed using a DigestPro robot (CEM) with the following protocol: washes with 25 mM ammonium bicarbonate and acetonitrile, reduction with 10 mM dithiothreitol at 60°C, alkylation with 50 mM iodoacetamide at room temperature, digestion with trypsin (Promega) at 37°C for 4 hours, and quenching with formic acid. The resulting peptides were analyzed by nano LC-MS/MS using a Waters M-class HPLC system interfaced with a ThermoFisher Fusion Lumos mass spectrometer. Peptides were loaded onto a trapping column and eluted over a 75 µm analytical column at 350 nL/min with a 30-minute gradient. MS and MS/MS spectra were acquired in the Orbitrap at resolutions of 60,000 and 15,000 FWHM, respectively. Data was searched using Mascot against the SwissProt Mouse database with a 10 ppm peptide mass tolerance and a 20 ppm fragment mass tolerance. Search parameters included trypsin digestion, fixed carbamidomethylation of cysteines, and variable modifications such as oxidation (M), acetylation (protein N-term), deamidation (NQ), and pyro-glu (N-term Q). Data was processed using Scaffold software with a 1% false discovery rate (FDR) at both the protein and peptide levels, requiring at least two unique peptides per protein.

### Gene ontology analysis

DAVID (Version 2021) (48) (https://david.ncifcrf.gov) was used to perform Gene Ontology analyses. Briefly, gene names were first converted to DAVID-recognizable IDs using Gene Accession Conversion Tool. The updated gene list was then subject to GO analysis using all Homo Sapiens as background and with Functional Annotation Chart function. GO_BP_DIRECT, KEGG_PATHWAY or UP_KW_BIOLOGICAL_PROCESS was used as GO categories. Only GO terms with a p value less than 0.05 were included for further analysis.

### Motif analysis

Motif analysis was performed with the SeqPos motif tool (version 0.590) (49) embedded in Galaxy Cistrome using all motifs within the mouse reference genome mm10 as background. LISA analysis was performed using webtool (http://lisa.cistrome.org/).

### Statistical Analysis

Data were analyzed and presented with GraphPad Prism software. Plots show individual data points and bars at the mean and ± the standard error of the mean (SEM). One-tailed t-tests were used to compare means between groups, with significance set at p < 0.05. In instances where the p value is not shown, *, **, ***, and **** represent p < 0.05, 0.01, 0.001, and 0.0001, respectively.

## Results

### Global RBBP5 binding to chromatin exhibits a ∼12h ultradian rhythm, concomitant with the promoter-proximal ∼12h H3K4me3, but not H3K9Ac/H3K27Ac rhythms in mouse liver

The 12h oscillator is transcriptionally regulated by the UPR TF XBP1s in mice (18,19,21), and likely under control by additional bZIP TFs, such as ATF4 and ATF6 (50). However, decades of research of eukaryotic gene regulation demonstrated that TFs themselves are not sufficient to drive transcription initiation and elongation, and additional so-called “co-regulators” are required for full gene activation by shaping the epigenetic landscape (51–53). Currently, the epigenetic landscape of both the 12h oscillator and proteotoxic stress responses are poorly characterized. To identify epigenetic regulators of global proteostasis control, we first examined a published temporal hepatic epigenome dataset for histone modifications associated with the active promoters of ∼12h genes (54). We found ∼12h rhythms of histone H3 trimethylated at lysine 4 (H3K4me3), but not two histone acetylation markers, histone H3 acetylated at lysine 9 (H3K9Ac) and lysine 27 (H3K27Ac), at the promoters of ∼12h genes (**Fig. 1A, B**). This is particularly prominent for those ∼12h genes with rhythmic expressions peaking at ZT0 and ZT12 that are enriched for proteostasis pathways (21), with H3K4me3 level also peaking at these two time points (**Fig. 1A, B**).

**Fig. 1.**
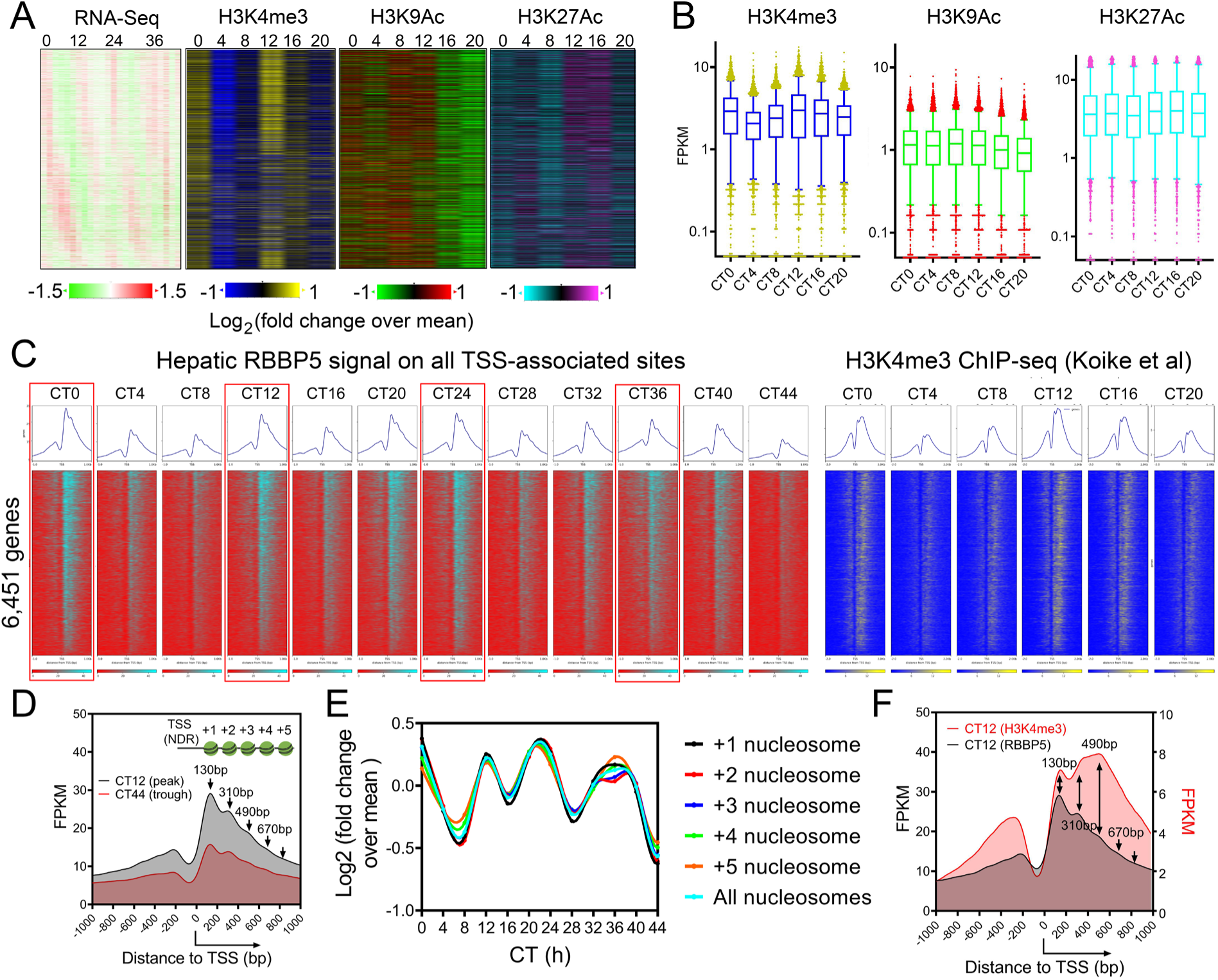
Global ∼12h RBBP5 cistrome is associated with promoter-proximal ∼12h H3K4me3 epigenome in mouse liver. **(A)** Heatmap of ∼12h hepatic transcriptome aligned with H3K4me3, H3K9Ac and H3K27Ac cistromes. **(B)** Quantification of temporal H3K4me3, H3K9ac and H3K27Ac. **(C)** Heatmap showing RBBP5 and H3K4me3 chromatin occupancy 1kb ± of TSS of 6,451 genes. **(D)** RBBP5 chromatin occupancy aligned with nucleosome positioning. **(E)** Quantification of temporal average RBBP5 binding intensity on different nucleosomes. **(F)** RBBP5 chromatin occupancy aligned with H3K4me3 occupancy.

In eukaryotes, H3K4 is tri-methylated by the COMPASS complex, which includes regulatory subunits RBBP5, ASH2L, WDR5, DPY30 and catalytic subunits SETD1A/B/MLL1-4 (**Fig. S1A**) (55–57). Via writing H3K4me3, COMPASS is essential for the eviction of paused RNA polymerase II (RNAPII) and promotes transcriptional elongation (58,59). Landscape In Silico deletion Analysis (LISA) (60) further inferred COMPASS subunits WDR5, ASH2L, DYP30 and RBBP5 as putative epigenetic regulators of hepatic ∼12h, but not circadian genes (**Fig. S1B**). Being the commonly shared regulatory subunits for all MLL family H3K4 methyl transferase complex (including SETD1A/B and MLL1-4), RBBP5/ASH2L forms the minimal heterodimer sufficient to activate all MLL histone methyl transferase (**61,62**) and are essential for the assembly of the whole COMPASS complex (**57,63**). Between the two essential subunits, we initially focused on RBBP5. Eigenvalue/pencil method (64) revealed that the levels of both hepatic *Rbbp5* mRNA and nuclear RBBP5 protein exhibit ∼12h oscillations, with the nuclear protein level peaking at ∼CT0 and ∼CT12 (**Fig. S1C, D**), in line with the acrophases of H3K4me3 and gene expression oscillations (**Fig. 1A, B**).

To further evaluate the potential role of RBBP5 in establishing the epigenetic landscape of hepatic ∼12h rhythms, we performed temporal RBBP5 ChIP-seq at a 4-h interval for a total of 48 hours in the liver of male C57BL/6 mice housed under constant darkness condition fed *ad libitum*. Chromatin occupancies of RBBP5 align well with nucleosome positioning downstream of the transcription start sites (TSS) and the H3K4me3 epigenome, with ‘ridges’ corresponding to 130bp, 310bp, 490bp, 670bp downstream of TSS at all time points, thus demonstrating the high quality of our ChIP-seq dataset (**Fig. 1C-F**). In total, we identified 7,963 hepatic RBBP5 binding sites that are within 2kb (up and downstream) of TSS of 6,451 genes (**Fig. 1C and Table S1**). The average binding intensities of RBBP5 on all post-TSS nucleosomes displayed robust 12h rhythms, with peaks occurring at CT0, CT12, CT24, and CT36 (**Fig. 1C-E**). These rhythms aligned with the 12h oscillations of promoter-proximal H3K4me3 (**Fig. 1C-E)**, but not with those of H3K9Ac or H3K27Ac (**Fig. S1E**). This pattern is exemplified by *Xbp1*, one of the major UPR TFs and the transcriptional regulator of the 12h oscillator, and *Hsph1*, a canonical 12h oscillator output gene encoding a heat shock protein (**Fig. S1F**). Using the eigenvalue/pencil and RAIN (40) algorithms, we identified 3,028 and 1,864 (p<0.05) RBBP5 binding sites that cycle with a ∼12h period, respectively (**Fig. S1G, H and Table S1**). Collectively, these results establish the epigenetic landscape of hepatic 12h oscillator is marked by RBBP5-H3K4me3.

### Hepatic RBBP5 cistrome coincides with that of XBP1s and hepatic ∼12h transcriptome

To investigate if RBBP5 regulates hepatic ∼12h rhythms of gene expression, we first examined the correlation between RBBP5 proximal promoter binding status and ∼12h rhythms of gene expression. As illustrated in **Fig. 2A**, genes with proximal promoter RBBP5 binding are enriched for ∼12h rhythms of gene expression (P=1.1e-39 by Chi-squared test). For those 2,525 ∼12h genes with RBBP5 binding, the phases of RBBP5 chromatin binding center around CT0 and CT12, slightly preceding the phases of gene expression observed at ∼CT1 and ∼CT13 (**Fig. 2B**). This temporal relationship suggests that RBBP5 acts as a driver, rather than a consequence of ∼12h rhythms of gene expression.

**Fig. 2.**
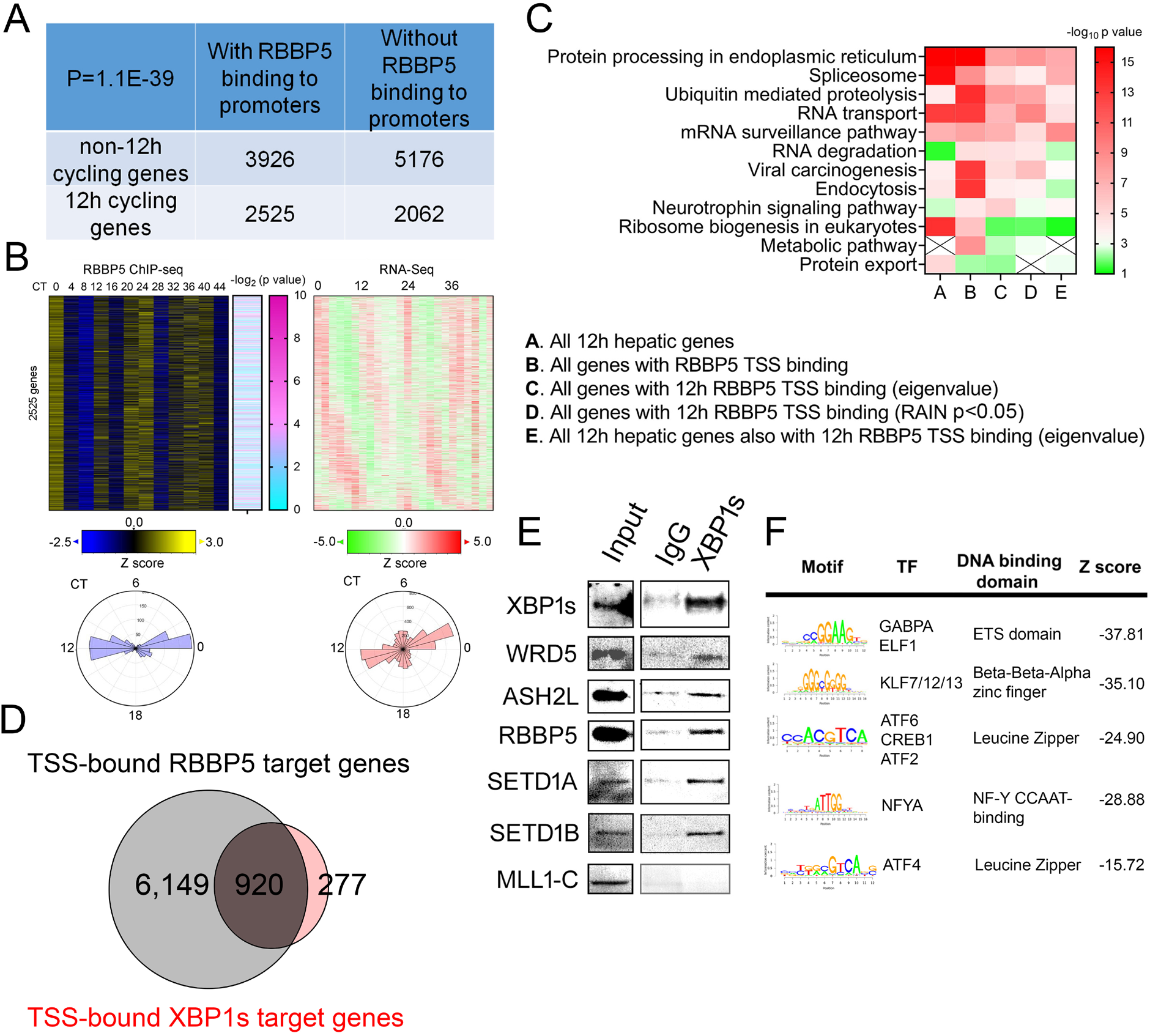
Hepatic RBBP5 cistrome coincides with that of XBP1s and hepatic ∼12h transcriptome. **(A)** The table shows the number of genes with or without ∼12h rhythms and with or without promoter RBBP5 binding. **(B)** Heatmap showing RBBP5 binding intensity along with - log_10_ transformed P values for having 1∼2h rhythm by RAIN and ∼12h rhythms of gene expression on 2,525 genes. **(C)** Heat map summary of GO analysis demonstrating the -log_10_ transformed P value of different enriched pathways for different genes. **(D)** Venn diagram demonstrating distinct and common cistromes for RBBP5 and XBP1s. **(E)** Western blot showing co-IP in liver nuclear extracts from CT0 using anti-XBP1s and normal IgG control antibody. **(F)** Motif analysis of RBBP5 binding sites of 6,149 genes.

We next performed Gene Ontology (GO) analysis on those hepatic genes with proximal promoter RBBP5 binding and revealed strong enrichment of proteostasis pathways including protein processing in the ER and ubiquitin-mediated proteolysis (**Fig. 2C**). Additionally, pathways involved in mRNA processing, such as spliceosome, RNA transport, and mRNA surveillance, were also enriched (**Fig. 2C**). These findings align with previous observations that both mRNA and protein metabolism pathways are enriched in genes exhibiting ∼12h rhythms in both mice and humans (16,17,65). The coupling of mRNA and protein metabolism is orchestrated by a SON-XBP1s axis and involves the liquid-liquid phase separation (LLPS) dynamics of nuclear speckles (17). Similar pathways are also enriched in genes exhibiting both ∼12h rhythms of RBBP5 promoter binding and gene expression (**Fig. 2C**).

XBP1s is the major transcriptional factor regulating the hepatic 12h oscillator and also exhibits a global 12h rhythm of chromatin binding to the promoter regions peaking at CT0, 12, 24 and 36 (19,21). Comparing the cistromes of RBBP5 and XBP1s revealed a significant overlap between the two, with 920 genes sharing RBBP5 and XBP1s proximal promoter binding (**Fig. 2D**). Co-immunoprecipitation in the hepatic nuclear extracts at CT0 confirmed the physical interaction between XBP1s and subunits of SET1/COMPASS, including RBBP5, ASH2L, WDR5, and two histone methyltransferase SETD1A and SETD1B (**Fig. 2E**). Motif analysis of the DNA sequences around the 6,149 RBBP5 binding sites that do not overlap with XBP1s cistrome identified binding motifs for GABPA, KLF, ATF6, NFYA and ATF4 (**Fig. 2F**). These findings suggest that, in addition to acting as a co-activator for XBP1s, RBBP5 may also work with other transcription factors to shape the hepatic 12h epigenome.

### RBBP5 is an epigenetic regulator of the ∼12h oscillator, but not the canonical ∼24h core circadian clock

To establish the causality between RBBP5 and hepatic 12h rhythms of gene expression, we generated RBBP5 liver hepatocyte-specific knockout (RBBP5 *^LKO^*) mice using the CRE-loxP system. Exon 10 of mouse *Rbbp5,* which encodes the SET/ASH2L binding domain, was flanked by loxP sites (**Fig. 3A**). CRE-mediated deletion of Exon 10 is expected to lead to a frameshift and nonsense-mediated RNA decay of truncated *Rbbp5* transcript (**Fig. S2A**). Crossing homozygous floxed *Rbbp5* mice with Albumin-CRE mice resulted in liver-specific deletion of RBBP5, which was confirmed by DNA genotyping, IF against RBBP5 and western blot analysis (**Figs. 3B and S2B-D**). The resulting RBBP5 *^LKO^* mice weighed slightly less than wild-type counterparts (**Fig. S3A**) but maintained normal rhythmic locomotor activity and fasting-feeding cycles under both 12h:12h L/D and constant darkness conditions (**Fig. S3B-I**).

**Fig. 3.**
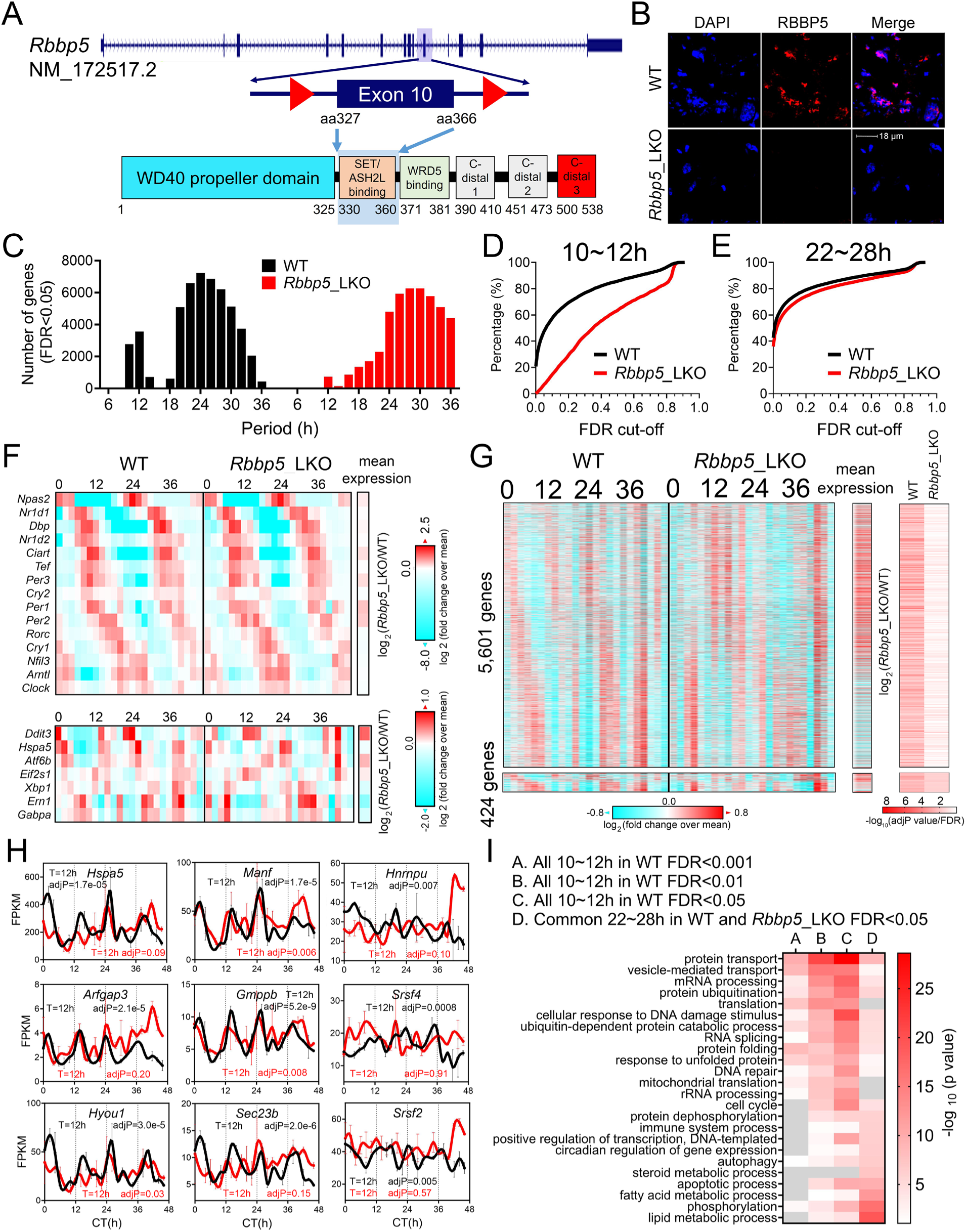
RBBP5 is an epigenetic regulator of the hepatic ∼12h oscillator, but not the canonical ∼24h circadian clock. **(A)** Schematic of the design of *Rbbp5^Flox^* mice. **(B)** Immunofluorescence of RBBP5 in the liver of *Rbbp5 ^Flox^* and *Rbbp5 ^LKO^* mice. **(C)** Histograms showing the period distributions of all rhythmic genes uncovered from *Rbbp5 ^Flox^* and *Rbbp5 ^LKO^* mice (with adjP values or FDR <0.05) (n=2 at each time point for either genotype). **(D, E)** Cumulative distribution of 10∼12h **(D)** and 22∼28h **(E)** genes in *Rbbp5 ^Flox^* and *Rbbp5 ^LKO^* mice with different FDR cut-offs. **(F)** Heat map of 15 core circadian clock (top) and canonical ∼12h gene expression (bottom) in the liver of *Rbbp5 ^Flox^* and *Rbbp5 ^LKO^* mice. Log_2_ normalized fold change of average gene expression across 48 hours between *Rbbp5 ^Flox^* and *Rbbp5 ^LKO^* mice was also shown. **(G)** Heat map of 6,025 ∼12h gene expression (FDR<0.05) in the liver of *Rbbp5 ^Flox^* and *Rbbp5 ^LKO^* mice, with 5,601 of them lost in *Rbbp5 ^LKO^* mice, along with -log_10_ transformed adjP values for having 10-12h rhythm by RAIN. Log_2_ normalized fold change of average gene expression across 48 hours between *Rbbp5 ^Flox^* and *Rbbp5 ^LKO^* mice was also shown. **(H)** RNA-Seq data for representative proteostasis and mRNA processing genes in the liver of *Rbbp5 ^Flox^* and *Rbbp5 ^LKO^* mice. **(I)** Heat map summary of GO analysis demonstrating the -log_10_ transformed P values of different enriched pathways for different genes. Data: Mean ± S.E.M.

To identify the RBBP5-dependent oscillating hepatic transcriptome, we performed bulk RNA sequencing (RNA-Seq) analysis in the liver of male *Rbbp5 ^Flox^* and *Rbbp5 ^LKO^* mice at 2-h intervals for a total of 48 hour under constant darkness in duplicates (**Table S2**). To identify genes cycling with a period ranging from 6 to 36 hours, we initially applied the RAIN algorithm (40) to each genotype’s temporal transcriptome (40) (**Table S3**). Compared to alternative methods like JTK_CYCLE, RAIN detects rhythms with arbitrary waveforms and therefore more robustly uncovers ultradian rhythms (21,23,39). Consistent with past studies (18,20,21,66,67), two populations of hepatic transcripts cycling with periods of ∼12h and ∼24h were identified in *Rbbp5 ^Flox^* mice liver (**Fig. 3C, D**). By contrast, in *Rbbp5 ^LKO^* mice, the ∼12h rhythms are all but abolished and the circadian oscillations remain largely intact, which include all core circadian clock genes (such as *Bmal1*, *Per1/2/3*, and *Nr1d1/2*) displaying robust circadian rhythms with the same phases and amplitudes as those observed in *Rbbp5 ^Flox^* mice (**Figs. 3C-F and S4A**). The maintenance of ∼24h circadian rhythms and abolishment of ∼12h ultradian rhythms in *Rbbp5 ^LKO^* mice is further confirmed by the principal component analysis (PCA) (**Fig. S4B, C**). Specifically, using RAIN with an FDR cutoff of 0.01 and 0.05, we identified a total of 3,549 and 6,025 hepatic transcripts cycling at periods between 10-12h in *Rbbp5 ^Flox^* mice, respectively. By contrast, only 61 and 813 10-12h genes were observed in *Rbbp5 ^LKO^* mice with the same FDR thresholds, indicating over 93% ∼12h hepatic transcriptome was abolished with liver RBBP5 ablation (**Figs. 3G, H, S4D and Table S3**). GO analysis confirmed that these RBBP5-depenent ∼12h hepatic transcriptomes are enriched in various mRNA processing and proteostasis pathways, distinct from those of RBBP5-independent circadian genes enriched in lipid metabolism and phosphorylation (**Fig. 3I**).

To determine the robustness of our results to different analytic methods, we also performed spectrum analysis with the eigenvalue/pencil method (**Table S4**) (19–21,23,26,39), which unlike statistical methods such as JTK_CYCLE and RAIN does not require pre-assignment of period range, enabling unbiased identification of multiple superimposed oscillations for any given gene (19–21,23,26,39). Revealing up to three oscillations from each gene, eigenvalue/pencil analyses also revealed prevalent ∼24h and ∼12h oscillations in *Rbbp5 ^Flox^* mice, along with a third population cycling at a period around 8h (**Fig. S5A-E**). Hepatic ∼8h oscillations were also previously reported but their regulation and function remained elusive (22,66). *Rbbp5 ^LKO^* mice, on the other hand, exhibited a drastically different spectrum, characterized by the near complete loss of the ∼12h rhythms, *de novo* gains of many faster ultradian rhythms with periods between 6 and 7h, and the maintenance of most circadian rhythms (**Fig. S5A-E**). Similar findings were also observed for dominant oscillations (which is the one with the largest amplitude among the three superimposed oscillations for each gene) for each gene (**Fig. S5A-E**). For the ∼12h genes originally identified in *Rbbp5 ^Flox^* mice but lost in *Rbbp5 ^LKO^* mice, the period spectrum in the *Rbbp5 ^LKO^* mice displayed a diverse distribution. This ranged from a complete lack of rhythmicity (∼39%) to shorter ultradian periods of 6-7 hours, and even to longer periods exceeding 24 hours (**Fig. S5F**). RBBP5-dependent ∼12h transcriptome uncovered by the eigenvalue/pencil showed substantial overlap with the results obtained through the RAIN method. This convergence was evident both in the specific genes identified (**Fig. S5C**) and in the enriched biological pathways (**Fig. S5G, H**), thereby confirming the robustness of our findings. These results thus establish RBBP5 as a central epigenetic regulator of the hepatic 12h oscillator, with more than 90% ∼12h hepatic transcriptome abolished in its absence.

In addition to liver, cell-autonomous ∼12h rhythms of proteostasis gene expression including *Xbp1* can also be observed in mouse embryonic fibroblasts (MEFs) synchronized by low concentration of ER stress inducer tunicamycin (Tu) (**21,23**). We observed a ∼12h rhythm of *Rbbp5* expression in Tu-synchronized MEFs, which is independent of BMAL1 but require XBP1s (**Fig. S6A**), indicating RBBP5 itself is also under 12h oscillator control. To test the functional role of COMPASS in regulating the cell-autonomous ∼12h rhythms, we knocked down either *Rbbp5* or *Ash2l* via siRNA in a previously described 12h oscillator reporter MEFs that stably express *Manf* promoter-driven destabilized luciferase (dluc) (21), and found that both siRNAs significantly dampen the ∼12h oscillation of luciferase activity as well as shorten the period (**Fig. S6B**). By contrast, neither *RBBP5* nor *ASH2L* knocking down affects the circadian oscillation of *Bmal1* promoter activity in human U2OS cells in a previous genome-wide siRNA screen study (68) (**Fig. S6C**). Collectively, our results demonstrate that RBBP5 is an integral component of the 12h oscillator, but dispensable for the canonical circadian clock both *in vivo* and *in vitro*.

### RBBP5 modulates the hepatic transcriptional response to acute proteotoxic stress *in vivo*

Before the discovery of the mammalian 12h oscillator, the dynamics of proteostasis were primarily studied as transient responses to various proteotoxic stresses, with a classic example being the UPR triggered by ER stress. In response to the accumulation of misfolded proteins in the ER, the UPR initiates a signaling cascade from the ER to the nucleus, ultimately activating three key transcription factors: XBP1s, ATF4, and ATF6. This activation leads to a beneficial adaptive response, upregulating proteostasis genes to restore ER homeostasis (**Fig. 4A**) (69–71). However, ER stress can also trigger maladaptive and deleterious responses, such as the activation of pro-inflammatory and immune gene expression through the TRAF2-mediated activation of AP1 (comprised of c-Fos and c-Jun), NF-κB or STAT3 TFs (**Fig. 4A**) (72). Our findings that RBBP5 can co-activate XBP1s to regulate the 12h oscillator suggest that RBBP5 is essential for the adaptive UPR TFs-mediated activation of proteostasis genes but is not required for the maladaptive activation of immune genes by immune-regulatory TFs, whose binding motifs are absent from RBBP5 binding sites (**Fig. 2H**).

**Fig. 4.**
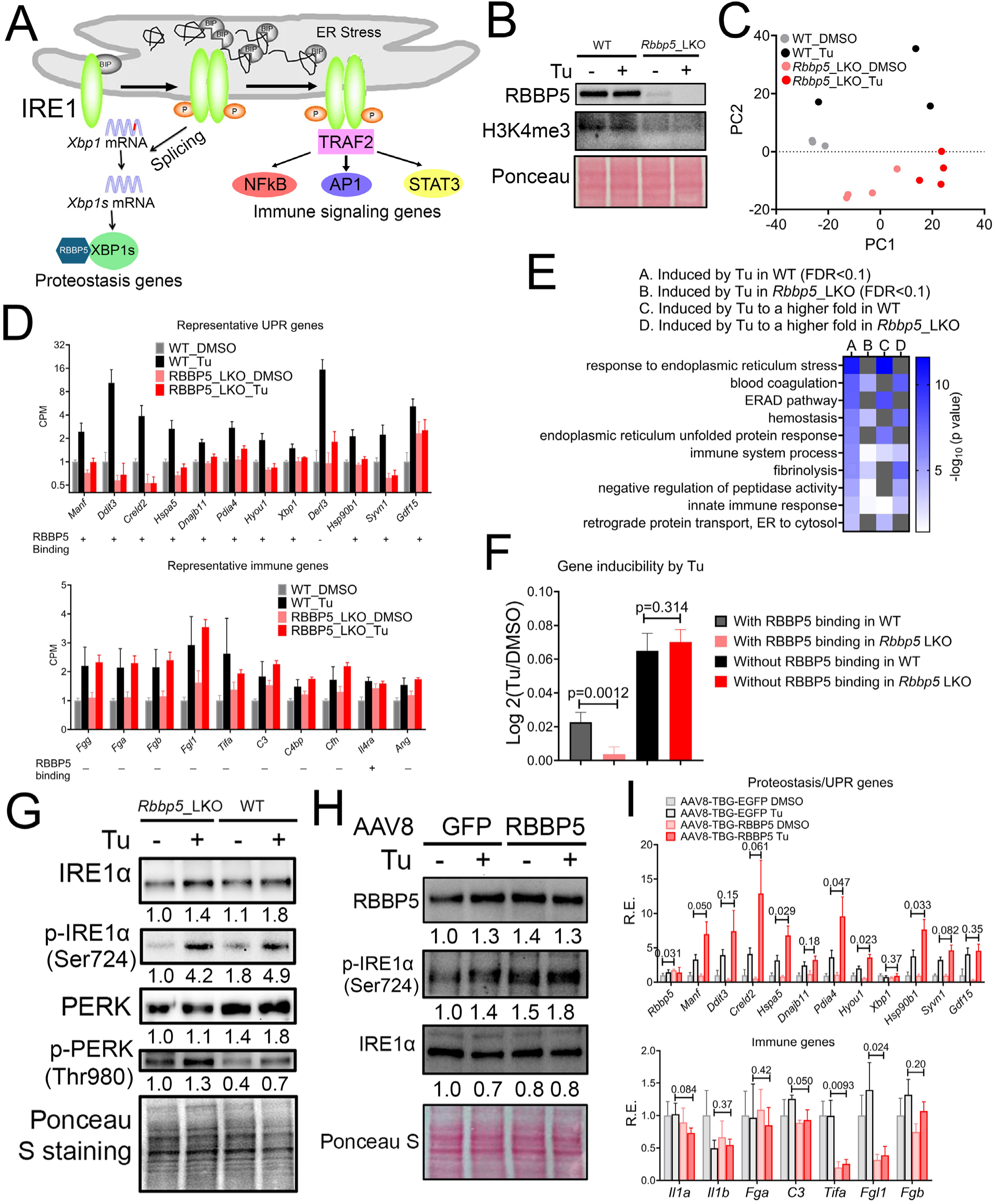
RBBP5 regulates the hepatic transcriptional response to proteotoxic stress. **(A)** The diagram of UPR. **(B)** Western blot analysis in the liver of *Rbbp5 ^Flox^* and *Rbbp5 ^LKO^* mice with or without Tu treatment. **(C)** PCA showing the hepatic transcriptional response to Tu in *Rbbp5 ^Flox^* and *Rbbp5 ^LKO^* mice. **(D)** Representative RNA-seq data for proteostasis (top) and immune genes (bottom) with RBBP5 promoter-binding status also shown below. n=3 for *Rbbp5 ^Flox^* DMSO and *Rbbp5 ^Flox^* Tu, and n=4 for *Rbbp5 ^LKO^* DMSO and *Rbbp5 ^LKO^* Tu. **(E)** Heat map summary of GO analysis demonstrating the -log_10_ transformed P values of different enriched pathways for different genes. **(F)** Log _2_ normalized fold induction by Tu in *Rbbp5 ^Flox^* and *Rbbp5 ^LKO^* mice for genes with or without RBBP5 promoter binding. **(G)** Western blot analysis of different proteins in *Rbbp5 ^Flox^* and *Rbbp5 ^LKO^* mice treated with or without Tu. **(H, I)** Mice tail-vein injected with AAV8-TGB-GFP or AAV8-TGB-RBBP5 were treated with or without Tu. Western blot analysis **(H)** and qPCR **(I)**. n=3 for AAV8-TBG-EGFP DMSO, n=4 for AAV8-TBG-EGFP Tu, n=4 for AAV8-TGB-mRBBP5 DMSO and n=4 for AAV8-TGB-mRBBP5 Tu. Data: Mean ± S.E.M.

To test this hypothesis, we intraperitoneally injected male *Rbbp5 ^Flox^* and *Rbbp5 ^LKO^* mice with vehicle control or 0.05mg/kg Tu to induce acute ER stress and performed RNA-seq on hepatic transcriptome isolated 8 hours post injection (**Table S5**). This lower dose of Tu was selected for its ability to induce the UPR, but that was significantly less toxic than the more common experimental dose of 1 mg/kg—which is known to elicit hepatocellular death and severe toxicity (73). Western blot analysis confirmed the depletion of RBBP5 and a reduction of H3K4me3 level in the liver of *Rbbp5 ^LKO^*mice (**Fig. 4B**). PCA demonstrated the separation of the four groups on the transcriptome space (**Fig. 4C**). Using an adjusted p value (FDR) of 0.10 as cut-off, we identified 139 and 45 genes significantly induced by Tu in the liver of *Rbbp5 ^Flox^* and *Rbbp5 ^LKO^*mice, respectively, with 21 commonly shared between the two (**Figs. 4D, S7A-G and Table S6**). In *Rbbp5 ^Flox^* mice, these 139 genes are enriched in pathways of proteostasis and immune functions as expected, with pathways of ‘response to ER stress’ and ‘ERAD’ among the top two GO terms (**Fig. 4D, E**). By contrast, in *Rbbp5 ^LKO^* mice, these 45 genes are only enriched in immune functions, with proteostasis genes not being significantly induced by Tu (**Fig. 4D, E**). In general, proteostasis genes were induced by Tu to a much greater fold in *Rbbp5 ^Flox^*mice (such as *Creld2*, *Xbp1*, *Hsp90b1*), while the opposite was found for blood coagulation and immune genes in *Rbbp5 ^LKO^*mice (such as *S100a9* and *Fgb*) (**Figs. 4D, E and S7E, F, H**). Further integrating RBBP5 ChIP-seq results, we found that, on average, the ablation of RBBP5 markedly diminished the fold induction by Tu for genes with RBBP5 promoter binding, while it had no effects for those without RBBP5 chromatin recruitment (**Fig. 4D, F**). To rule out the possibility that RBBP5 regulates UPR via influencing the upstream ER stressing in the ER, we performed western blot analysis of p-IRE1α and p-PERK in the liver of *Rbbp5 ^Flox^* and *Rbbp5 ^LKO^* mice and did not observe any notable difference in the increase of their phosphorylated level in response to Tu between the two genotypes (**Fig. 4G**).

To determine if RBBP5 overexpression is also sufficient to amplify UPR, we tail-vein injected AAV8-TBG-GFP and AAV8-TBG-RBBP5 viruses into wild-type male mice followed by intraperitoneal injection with vehicle control or 0.025mg/kg Tu 30 days later. Thyroxine binding globulin (TBG) promoter is liver-specific and directs RBBP5 transgene expression in hepatocytes (74). qPCR and western blot analysis confirmed an increase of total hepatic expression of RBBP5 of ∼1.4 fold in AAV8-TBG-RBBP5 group under vehicle control condition (**Fig. 4H, I**). Consequently, RBBP5 hepatic overexpression significantly amplified the UPR of proteostasis genes including *Manf*, *Hspa5* (*BiP*), *Pdia4*, *Hyou1*, and *Syvn1* without affecting the upstream ER stress sensing (**Fig. 4H, I**). In addition, we found that RBBP5 overexpression attenuated the expression of several immune and blood coagulation genes such as *C3*, *Tifa*, *Fgl1* and *Fgb* under ER stress conditions (**Fig. 4I**). Taken together, these results demonstrate that RBBP5 is both necessary and sufficient for the transcriptional regulation of hepatic response to acute proteotoxic stress *in vivo*. RBBP5 activates the expression of proteostasis genes while repressing those involved in immune functions.

### RBBP5 protects mice from chronic ER stress-induced hepatic inflammation and steatosis

Contrary to acute ER stress, chronic ER stress, experimentally reconstituted by repeated injection of Tu, paradoxically leads to feedback-mediated suppression of the UPR signaling, to even below basal levels (73). Proposed mechanisms of this suppression include silencing of the ATF6a branch of UPR, enhancement of mRNA degradation via IRE1-dependent decay (RIDD), and the direct inhibition of UPR stress sensors by the ER chaperone HSPA5 (BiP) (73). To investigate whether RBBP5 also modulates the feedback suppression of UPR under chronic ER stress conditions, we repeatedly intraperitoneally injected male *Rbbp5 ^Flox^* and *Rbbp5 ^LKO^*mice with vehicle control or 0.025mg/kg Tu for six consecutive days and harvested liver tissues 24 hours after the last injection for analysis (**Fig. 5A**). Western blot analysis revealed attenuation of p-IRE1α and ATF4 level in both *Rbbp5 ^Flox^* and *Rbbp5 ^LKO^* mice with repeated Tu injection (**Fig. 5B**). In agreement with the Western blot result, qPCR analysis also confirmed the downregulation of proteostasis genes *Atf4*, *Hyou1*, and *Manf* in both mice under chronic ER stress condition (**Fig. 5C**). By contrast, compared to *Rbbp5 ^Flox^* mice, the expression of pro-inflammatory cytokine genes *Il1α* and *Il1b* were higher in Tu-injected *Rbbp5 ^LKO^* mice, consistent with higher levels of p-IRE1α and p-c-Jun (Ser73) observed in *Rbbp5 ^LKO^* mice (**Fig. 5B, C**). The persistent elevated levels of p-IRE1α/p-c-JUN and pro-inflammatory cytokine gene expression in *Rbbp5 ^LKO^* mice suggest that these mice exhibit heightened susceptability to Tu-induced ER stress and inflammation. In summary, our results indicate that while wild-type mice can sustain persistent cycles of UPR activation and deactivation in response to ER stress, as previously described (73), the loss of RBBP5 disrupts this balance. Although RBBP5-deficient mice retain the ability to attenuate UPR, they are unable to properly activate it. Consequently, *Rbbp5 ^LKO^* mice exhibit consistently lower expression of adaptive proteostasis genes under ER stress condition, leading to heightened proteotoxic stress and chronic inflammation likely via activating the p-IRE1α-TRAF2-AP1 signaling cascade.

**Fig. 5.**
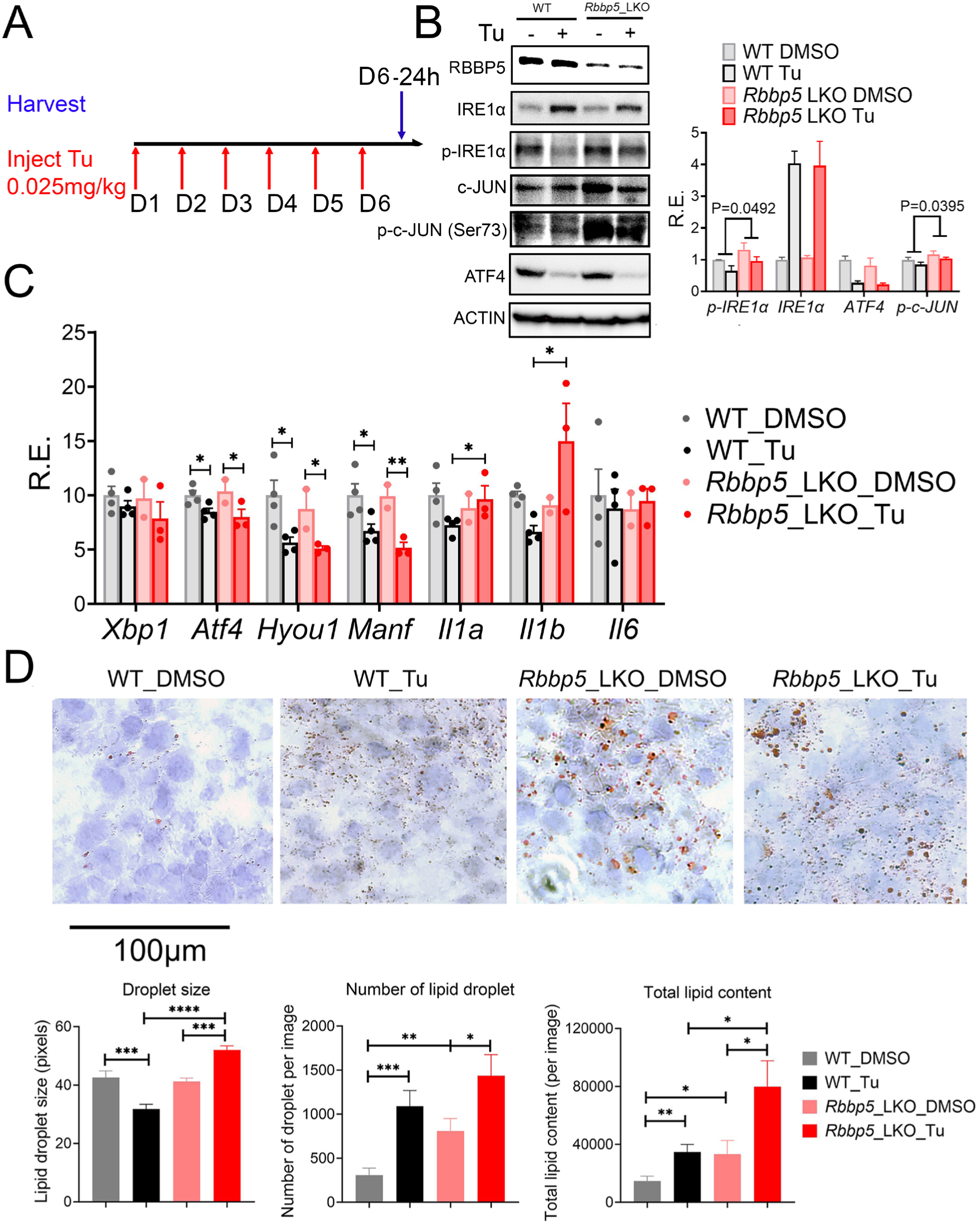
RBBP5 protects mice from chronic ER stress-induced hepatic inflammation and steatosis. **(A)** A schematic of experimental design. **(B)** Western blot and quantification of the expression of different proteins in *Rbbp5 ^Flox^* and *Rbbp5 ^LKO^* mice with or without repeated Tu injection. **(C)** qPCR analysis of different proteostasis and immune gene expressions in the same cohort of mice. **(D)** Representative images of Oil Red O staining and quantification of lipid droplet size, number and total lipid content (total lipid droplet area). n=4 for *Rbbp5 ^Flox^* DMSO, n=4 for *Rbbp5 ^Flox^* Tu, n=2 for *Rbbp5 ^LKO^* DMSO, n=3 for *Rbbp5 ^LKO^* Tu. Data: Mean ± S.E.M.

Both chronic ER stress and inflammation can result in hepatic steatosis (75–78). We therefore performed Oil Red O staining, which stained all neutral lipids, on liver tissues from *Rbbp5 ^Flox^* and *Rbbp5 ^LKO^* mice after repeated vehicle or Tu injection. *Rbbp5 ^LKO^* mice liver displayed an increased number of lipid droplets and total lipid content, compared to wild-type counterparts under both basal and chronic ER stress conditions (**Fig. 5D**). Our data revealed that RBBP5 protects mice from chronic ER stress-induced hepatic inflammation and steatosis.

### Loss of RBBP5 compromises the cell-autonomous transcriptional response to proteotoxic stress and nutrient-deprivation, resulting in impaired autophagy and reduced cell survival

To determine whether RBBP5 regulates transcriptional responses to proteotoxic stress in a cell-autonomous manner beyond the liver, we examined MEFs, which exhibit both a robust cell-autonomous XBP1s-dependent 12h oscillator and strong transcriptional responses to proteotoxic stress (17,18,21,79). *Rbbp5* mRNA expression was induced by Tu (**Fig. 6A**), and transient knockdown of RBBP5 using siRNA (**Fig. S8A**) significantly dampened the UPR of proteostasis gene expressions in response to Tu, mirroring the effects observed with XBP1s knockdown (**Fig.6A**). Similarly, knocking down ASH2L also markedly dampened the UPR, whereas knocking down SETD1A, one of the H3K4me3 methyltransferases in the SET1/COMPASS complex, had only modest effects, which were likely due to redundancy among the many H3K4me3 methyltransferases that exist (**Fig. S8B**). As a negative control, we further knocked down both GCN5 and PCAF, proteins that function to write H3K9Ac as subunits in two distinct histone acetyltransferase (HAT) complexes, SAGA (Spt-Ada-Gcn5 acetyltransferase) and ATAC (Ada2a-containing) (80–82) and found it had no effects on the induction of *Xbp1* and *Manf* expression in response to ER stress (**Fig. 6B**). The lack of effects on UPR for HAT subunits is consistent with the absence of ∼12h rhythms of both H3K9Ac and H3K27Ac levels around the promoter of ∼12h proteostasis genes in mouse liver (**Fig. 1A, B**). ChIP-qPCR analysis confirmed a gradual, temporal increase in the recruitment of RBBP5 to the proximal promoters of select proteostasis genes in response to ER stress, concomitant with an increase in promoter-proximal H3K4me3 levels (**Fig. 6C**). Since XBP1s is the major TF driving UPR, we further performed IF on XBP1s and RBBP5 in MEFs and found a strong recruitment of RBBP5 to XBP1s in response to ER stress (**Fig. 6D, E**).

**Fig. 6.**
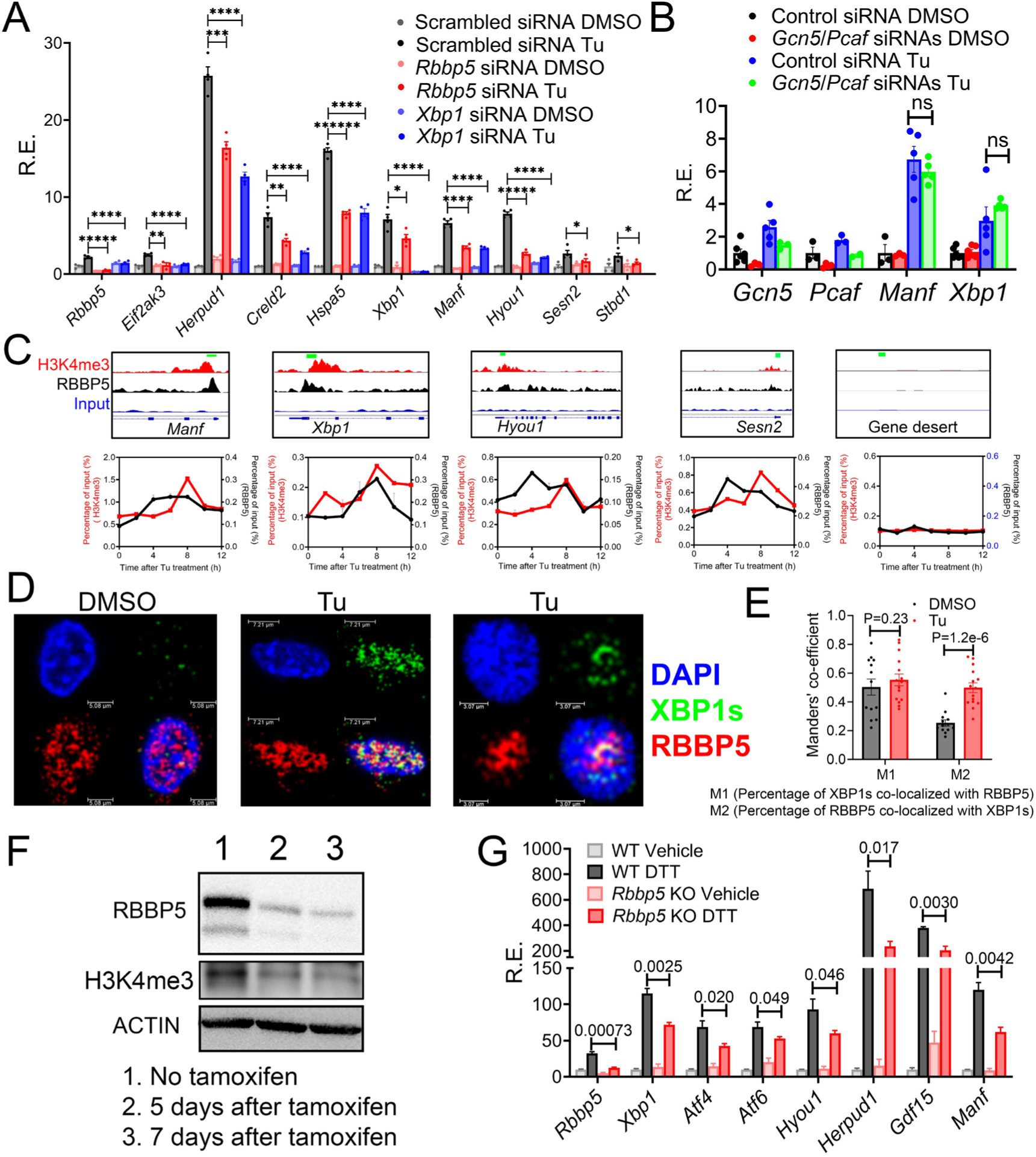
RBBP5 is required for cell-autonomous transcriptional response to ER stress. **(A)** qPCR of different genes in MEFs transfected with scrambled, *Rbbp5* or *Xbp1* siRNAs and treated with DMSO or Tu (100ng/ml) for 8 hours. **(B)** qPCR of different genes in MEFs transfected with scrambled, or the combination *Gcn5* and *Pcaf* siRNAs and treated with DMSO or Tu (100ng/ml) for 8 hours. (**C**) Selected genes aligned for RBBP5 and H3K4me3 ChIP-seq signal from CT0 mouse liver and ChIP-qPCR of RBBP5 and H3K4me3 in MEFs treated with Tu for different hours. (**D, E**) Immunofluorescence of XBP1s and RBBP5 in MEFs treated with vehicle control (DMSO) or Tu (100ng/ml) for 8 hours (**D**) and quantification of Manders’ coefficient of colocalization of XBP1s with RBBP5 (**E**). (**F**) *Rbbp5* (fl/fl) ROSA26-CreERT2 (+/+) MEFs were treated with tamoxifen for 5 or 7 days and western blot analysis was performed. (**G**) qPCR of different genes in *Rbbp5* (fl/fl) ROSA26-CreERT2 (+/+) MEFs treated with vehicle (WT) or tamoxifen (*Rbbp5* KO) followed by treatment with vehicle control or 1mM DTT for 5 hours. Data: Mean ± S.E.M.

To broadly assess the function of RBBP5 in the transcription regulation of proteotoxic stress response, we utilized orthogonal approaches to both deplete RBBP5 expression and trigger cellular stress. We took advantage of our homozygous *Rbbp5 ^Flox^* mice by crossing them with homozygous ROSA26 Cre-ERT2 mice, then isolated and immortalized primary MEFs from homozygous F2 embryos (**Fig. S8C**). Treating these *Rbbp5* (flox/flox); ROSA26 Cre-ERT2 (+/+) MEFs with tamoxifen for 7 days resulted in the complete deletion of both floxed *Rbbp5* alleles (**Fig. S8D**) and a subsequent ∼90% reduction in RBBP5 protein and a reduction of H3K4me3 level (**Fig. 6F**). The remaining RBBP5 protein is likely extremely long-lived and remains stable even after the deletion of underlying coding genes. To trigger ER stress, we used a different stimulus: Dithiothreitol (DTT), which disrupts protein disulfide formation in the ER (83,84). Similar to Tu, RBBP5 depletion significantly dampened the induction of proteostasis gene expression in response to DTT (**Fig. 6G**).

Next, we tested two additional proteotoxic stressors that are more physiologically relevant: leucine deprivation to simulate amino acid restriction, and a more severe nutrient deprivation stress by culturing cells in Earle’s Balanced Salt Solution (EBSS) for 16 hours. Both treatments not only trigger UPR but also activate the broader integrated stress response (ISR), resulting in the induction of autophagy (85). RBBP5 depletion impaired the transcriptional response to both leucine deprivation and EBSS, leading to a reduced induction of autophagy-promoting genes (**Fig. 7A**). Transcriptome profiling by RNA-seq confirmed the globally defective adaptive stress response to nutrient deprivation (EBSS) in the absence of RBBP5, where only 327 out of 2,351 genes can still be induced (**Figs. 7B-D, S8E, and Table S7, 8**). To investigate the physiological consequence of RBBP5 deletion, we quantified the autophagic flux by utilizing a tandem LC3 reporter mCherry-GFP-LC3 where an increase in the number of red-fluorescent cytosolic puncta indicates increased autolysosome formation and autophagic flux (**Fig. 7E**) (85). Consistent with the qPCR results, RBBP5 depleted MEFs failed to induce a robust general autophagy in response to leucine deprivation and further exhibited lower basal level of autophagic flux (**Figs. 7F and S8F**). Western blot analysis of p62/SQSTM1, a marker of autophagic flux due to its degradation during autophagy (86), showed a significant reduction in wild-type, but not RBBP5 depleted cells following leucine deprivation, thus confirming the defective autophagy phenotype associated with RBBP5 depletion (**Fig. S8G, H**). Since prolonged nutrient deprivation, such as 16h of EBSS treatment, can robustly induce both general autophagy and autophagy of the ER (ER-phagy) (87), we further measured ER-phagy by utilizing a previously published ER-phagy reporter system where a stably expressed ER-targeting mCherry-RAMP4 fusion is cleaved to free mCherry during starvation-induced ER-phagy (29) (**Fig. 7G**). As shown in **Fig. 7H, I**, RBBP5 depletion prevented the cleavage of mCherry in response to EBSS, indicating an impaired ER-phagy. Western blot analysis of p62/SQSTM1 further showed undetectable levels in wild-type cells following EBSS treatment, whereas a significant level remained in RBBP5 depleted cells under the same conditions, thus confirming a reduction of the general autophagy in RBBP5 depleted cells in response to EBSS (**Fig. 7H**). Consistent with weakened stress response and impaired autophagy, RBBP5 depleted MEFs displayed reduced cell survival under nutritional stress (**Fig. S8I**). Taken together, our results demonstrate that RBBP5 is indispensable for regulating the cell-autonomous transcriptional responses to various proteotoxic stresses as a core subunit of the H3K4me3-writing COMPASS complex.

**Fig. 7.**
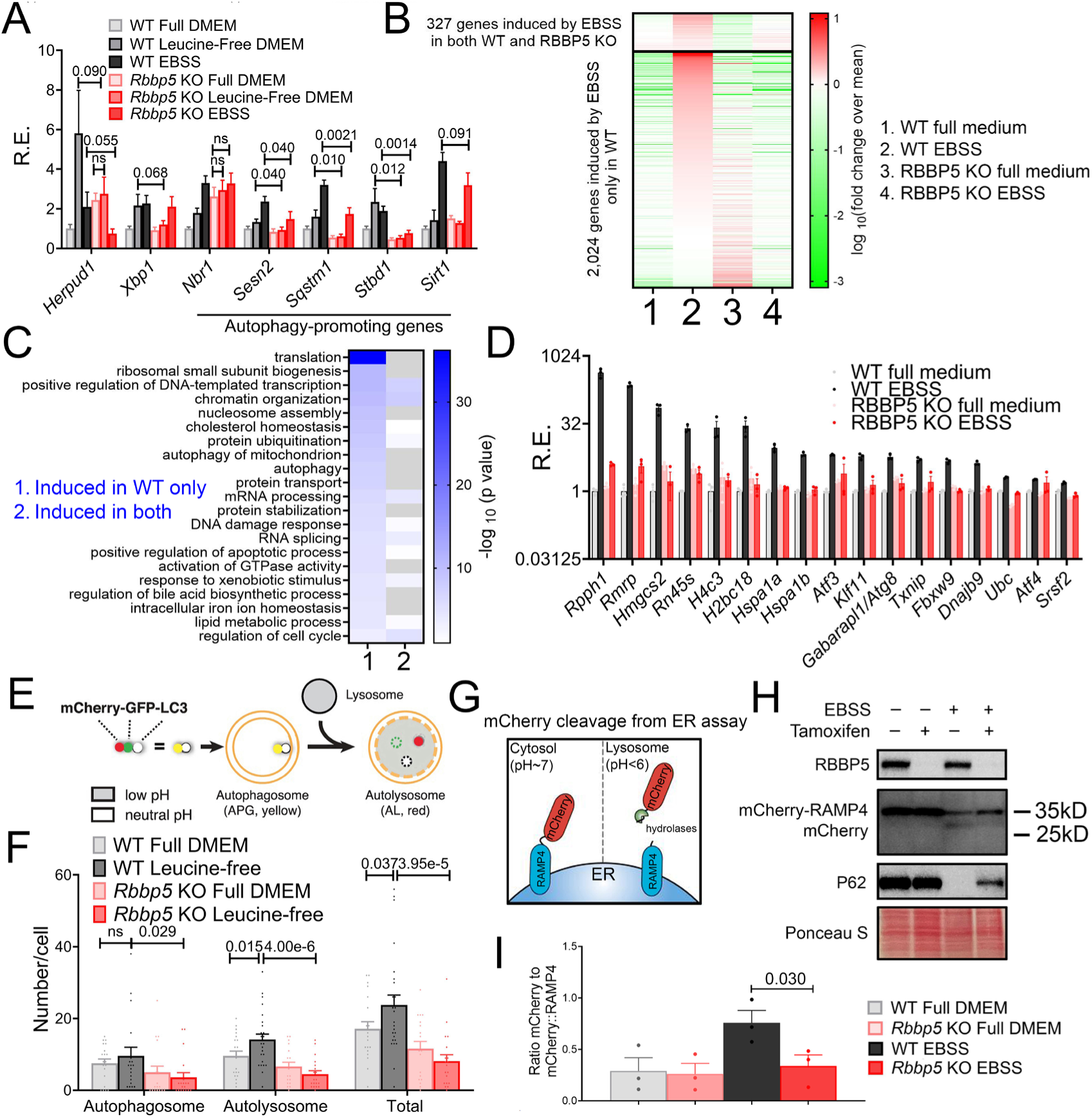
RBBP5 is required for cell-autonomous transcriptional response to nutrient deprivation. **(A)** qPCR analysis of *Rbbp5* (fl/fl) ROSA26-CreERT2 (+/+) MEFs pre-treated with (RBBP5 KO) or without (WT) tamoxifen and then treated with full DMEM media, leucine-free media or EBSS for 16 hours. **(B-D)**. Transcriptome profiling of *Rbbp5* (fl/fl) ROSA26-CreERT2 (+/+) MEFs pre-treated with (RBBP5 KO) or without (WT) tamoxifen and then treated with full DMEM media or EBSS for 16 hours by RNA-seq. Heatmap of all genes that are induced by EBSS in WT cells by EBSS (adjP<0.05) (**B**). GO analysis of EBSS-induced genes in only WT or both genotypes (**C**), and representative gene expression (**D**). **(E)** Diagram of the mCherry-GFP-LC3 autophagy reporter. **(F)** Quantification of autophagic flux in mCherry-GFP-LC3-expressing *Rbbp5* (fl/fl) ROSA26-CreERT2 (+/+) MEFs treated with vehicle (WT) or tamoxifen (*Rbbp5* KO) followed by treatment with full DMEM media or leucine-free media for 16 hours. **(G)** Diagram of the ER-phagy reporter using the mCherry cleavage from ER assay. **(H, I)** Western blot of different proteins **(H)** and quantification **(I)** of mCherry cleavage in RAMP4-mCherry-expressing *Rbbp5* (fl/fl) ROSA26-CreERT2 (+/+) MEFs treated with vehicle (WT) or tamoxifen (*Rbbp5* KO) followed by treatment with full DMEM media or EBSS for 16 hours. Data: Mean ± S.E.M.

### Proximity labeling reveals H3K4me-associated proteomic architecture of the proteostatic stress response

H3K4me3 is widely recognized as an active promoter mark, but its precise role in transcriptional regulation remains a subject of debate. Initially, it was believed that H3K4me3 facilitated transcription initiation by recruiting the RNAPII preinitiation complex (88), However, recent evidence suggests that H3K4me3 primarily regulates RNAPII pause-release and contributes to transcriptional elongation, likely through the recruitment of Integrator Complex subunit 11 (INTS11) (58,59). This discrepancy may stem from the context-specific functions of H3K4me3 in recruiting different transcriptional regulators across different biological processes (89). To shed light on the potential mechanisms by which H3K4me3 modulates proteostasis stress response, we performed quantitative TurboID-based proximity labeling in MEFs (in triplicates per condition), utilizing anti-H3K4me3 antibody (with anti-IgG as negative control) followed by recombinant protein A-TurboID labeling in isolated nuclei of MEFs before (T=0h) and 2.5h and 8h after ER stress induction by Tu (**Figs. 8A, S9A**) (47). While the total level of H3K4me3 remains largely unchanged during ER stress (**Fig. 8B**), we identified a total of 239 proteins (p<0.1) whose recruitment to H3K4me3 peaked at 8h post Tu treatment (**Table S9**). Pathway analysis of these 239 proteins revealed expected enrichment of COMPASS subunits (**Figs. 8C, S9B, C**), RNAPII subunits (**Fig. S9D**), as well as previously identified Integrator Complex subunits (such as INTS11) (**Fig. S9E**). The previously identified H3K4me3 demethylase KDM5A (59) revealed a much more constant recruitment to H3K4me3 during ER stress, with only a slight increase at 8h after ER stress induction (**Fig. S9C**).

**Fig. 8.**
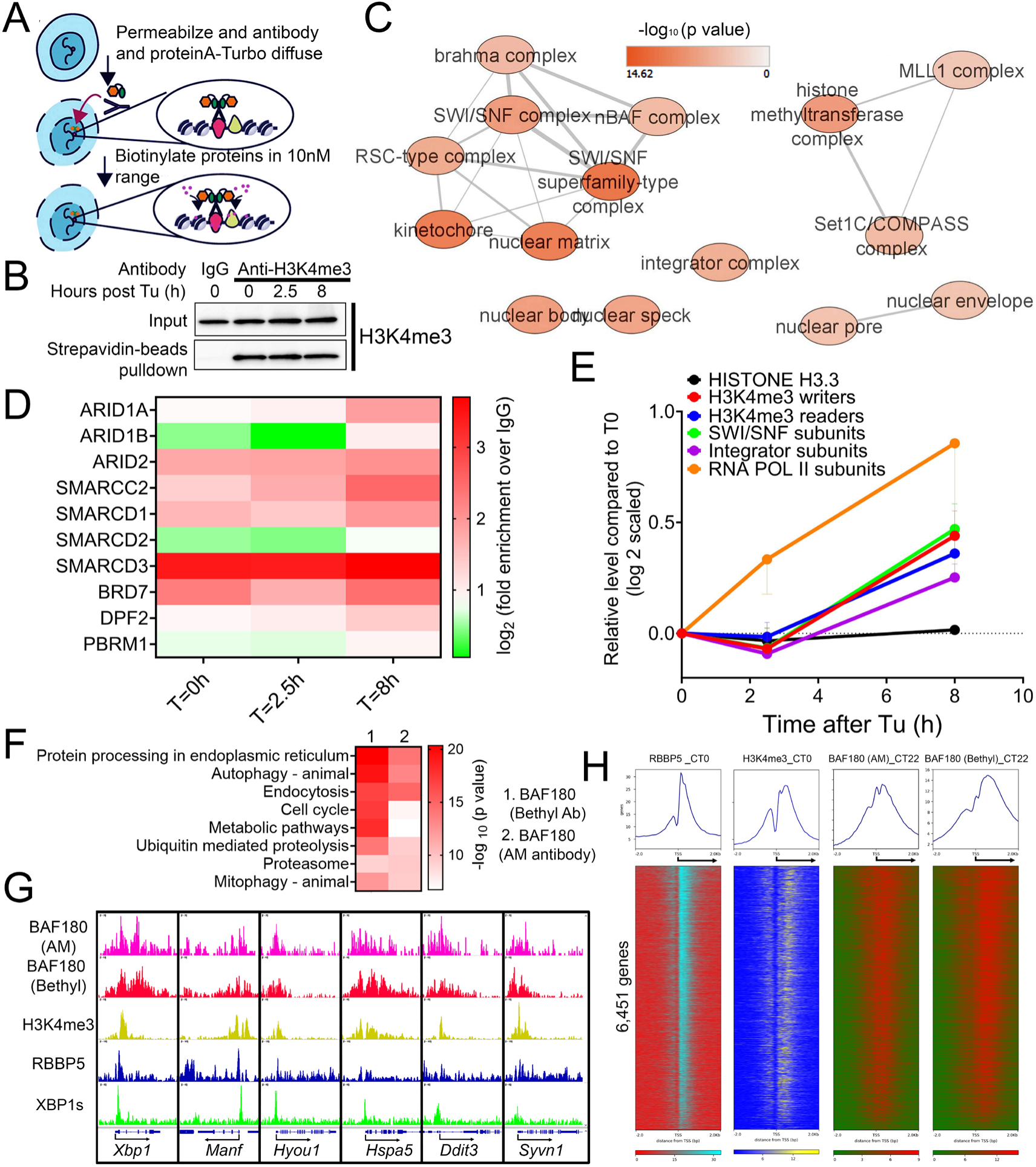
Proximity labeling reveals H3K4me-associated proteomic architecture of the proteostatic stress response. (**A**) A diagram illustrating the procedure of performing H3K4me3 proximity labeling via using the recombinant Protein A-TurboID. (**B**) Western blot analysis showing the nuclear level of total (input) or biotinylated (after streptavidin beads pull-down) level of H3K4me3 in MEF treated with 100ng/ml Tu for 0, 2.5 or 8h before subject to anti-IgG and anti-H3K4me3 antibody-based proximity labeling. (**C**) Enriched biological network of 239 proteins. (**D**) Heatmap of fold enrichment of different SWI/SNF subunits (over IgG control) at different times after ER stress. (**E**) Quantification of the average relative enrichment of proteins in different complexes at different times after ER stress. (**F**) GO analysis of BAF180 hepatic cistrome. (**G**) Selected UPR genes aligned for BAF180, RBBP5, H3K4me3 and XBP1s ChIP-seq signal in mouse liver. (**H**) Heatmap showing RBBP5, H3K4me3 and BAF180 chromatin occupancy 2kb ± of TSS of 6,451 genes. Data: Mean ± S.E.M.

Besides previously identified H3K4me3 writer/eraser/readers, we also revealed strong enrichment of SWI/SNF chromatin remodelers on H3K4me3 peaking at 8h after ER stress induction (**Fig. 8C, D**). These include BAF-specific (ARID1A/B), PBAF-specific (ARID2, PBRM1 and BRD7) and shared subunits (SMARCD1/2/3 and SMARCC2) (**Fig. 8C, D**) (90). The timing of SWI/SNF recruitment to H3K4me3 during ER stress closely parallels that of COMPASS and Integrator subunits but lags behind RNAPII subunits (**Fig. 8E**). This temporal pattern suggests that SWI/SNF might potentially play a role in facilitating transcriptional elongation during stress-induced gene activation.

To uncover further evidence supporting the recruitment of SWI/SNF to H3K4me3 during proteostatic stress, we turned back to the hepatic 12h oscillator and examined the temporal expression and cistrome of SWI/SNF subunits in mouse liver. Intriguingly, even before our discovery of the hepatic 12h oscillator in 2017, we had previously reported the 12h rhythmic expression of nuclear PBRM1 (BAF180), ARID2 (BAF200), and SMARCA4 (BRG1) proteins in wild-type mouse liver (36), though at the time, we were unaware of their potential implications in proteostasis control. To validate the hepatic 12h expression of nuclear PBRM1 (BAF180), we performed western blot analysis and confirmed its rhythmic peaks at CT10 and CT22 (**Fig. S10A**). Next, we conducted a post-hoc analysis of our previously published hepatic BAF180 ChIP-seq datasets at CT22, using two different antibodies (Bethyl and Active Motif) (35), and compared them with RBBP5 and H3K4me3 cistromes. Globally, genes with promoter-proximal BAF180 binding (±1 kb from the TSS) were significantly enriched in proteostasis-related pathways, including ER stress, autophagy, and proteolysis (**Fig. 8F, G**). Concordantly, we observed co-enrichment of BAF180, RBBP5, and H3K4me3 at the promoter regions of the 6,451 genes identified in **Fig. 1C**, with a strong positive correlation between their binding intensities on these genes (**Figs. 8H, S10B, C**). Although the causal relationship between SWI/SNF subunits and proteostasis gene expression remains to be determined, our collective findings in both MEFs and liver establish both COMPASS and SWI/SNF as critical components of the chromatin landscape governing dynamic proteostasis gene regulation.

### Reduced RBBP5 expression is associated with aging in mice and hepatic steatosis in humans

The roles of RBBP5 in diseases are incompletely understood (56,91). Since the decline of proteostasis and stress response is causally associated with both aging and metabolic diseases (92–95), we aim to uncover potential implications of RBBP5 in these two processes. Intriguingly, in a recent study, RBBP5 (and DPY30) is predicted to be a positive regulator of maximum lifespan across 26 different species, with its major effect predicted to be related to liver (96). Consistent with this prediction, an overall reduction of *Rbbp5* mRNA level is observed in the liver of aging mice (97), largely concordant with the expression dynamics of 97 stress response genes (those genes exhibiting 12h expression, directly bound by RBBP5 and can be induced by ER stress) (**Fig. 9A**).

**Fig. 9.**
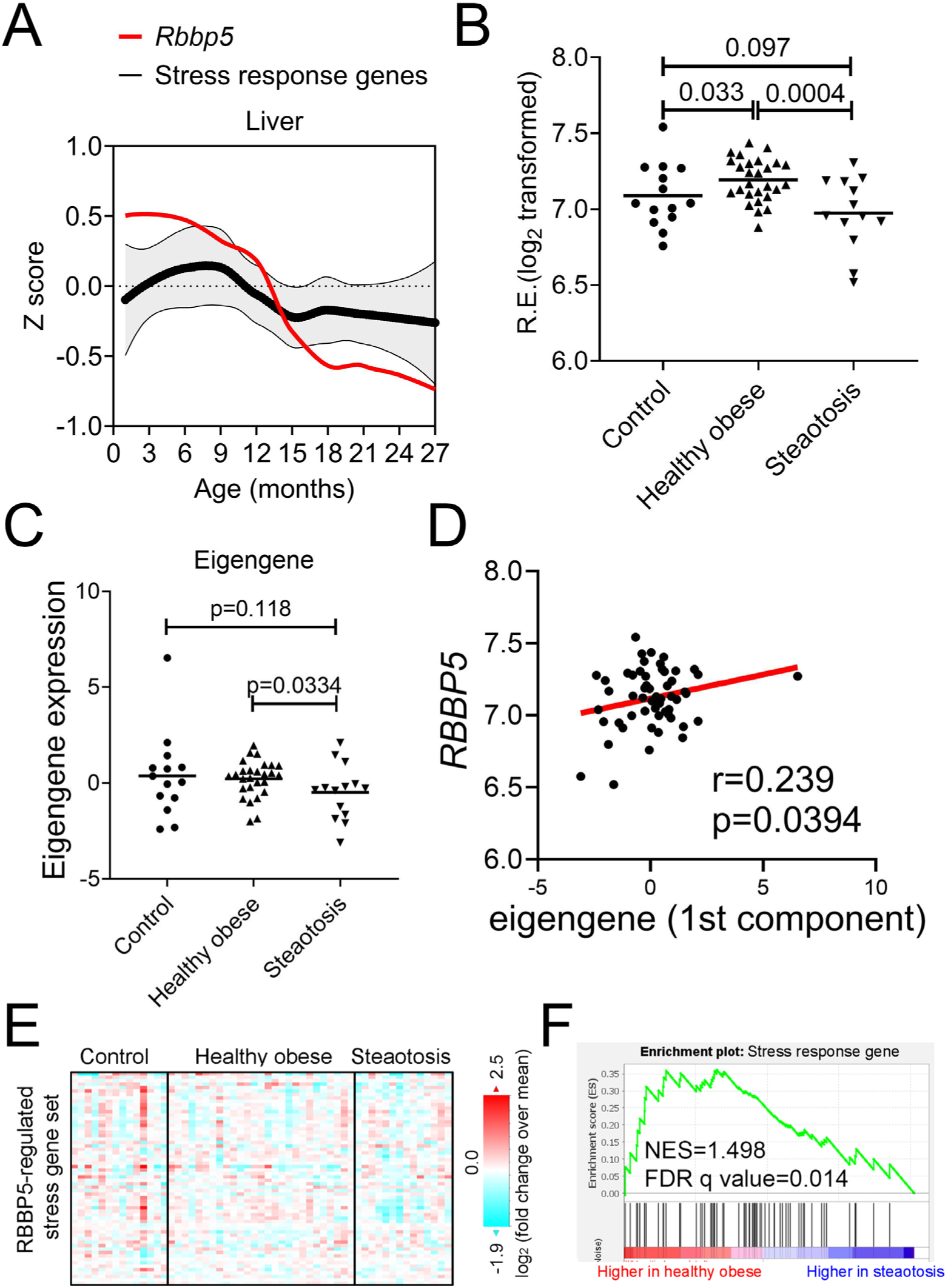
Reduced RBBP5 expression is associated with aging in mice and hepatic steatosis in humans. **(A)** Z score-normalized expression of mouse *Rbbp5* and average expression of stress response genes across mouse life span according to Tabua Muris dataset (97). **(B-F)** Data in humans with NAFLD/MAFLD was from (99). Expression of *RBBP5* (**B**) and the eigengene of 97 stress genes (**C**). (**D**) Scatter plot showing a positive correlation between *RBBP5* and eigengene expression. **(E)** Heatmap of 97 stress genes expression in different cohorts of human subjects. (**F**) GSEA analysis showing stress gene expression are downregulated in steatosis subjects.

Leveraging human gene-derived correlations across tissues (GD-CAT) dataset (98), we identified in human liver, RBBP5 expression is positively correlated with those involved in proteostasis (such as ribosome biogenesis, translation regulation) and mRNA metabolism (mRNA splicing and processing) and negatively associated with those implicated in immune response (acute phase response, complement activity, and immunoglobin activity) (**Fig. S11**). These results are in line with our RNA-seq and ChIP-seq data obtained from *Rbbp5 ^Flox^* and *Rbbp5 ^LKO^* mice (**Figs. 2C, 3I, 4D**). In humans with NAFLD/MAFLD (99), compared to healthy obese, RBBP5 mRNA level is significantly reduced in steatosis subjects (**Fig. 9B**), concordant with reduced stress gene expression in the same cohorts (**Fig. 9C-F**). Overall, a positive correlation between *Rbbp5* and the eigengene expression of stress response genes [eigengene (100) is the principle component of 97 stress response genes expression in each subject] is observed across all subjects (**Fig. 9D**). Collectively, these data suggest that the downregulation of *Rbbp5* and proteostatic stress response genes expression is associated with aging and MAFLD in mice and humans, respectively.

## Discussion

While the epigenetic landscape of the mammalian ∼24h circadian clock was well-studied (54,101–104), the epigenome of the mammalian ∼12h ultradian oscillator remains completely unexplored. In this study, we uncovered that unlike the circadian clock, which was predominantly regulated by histone acetyltransferase (HAT) (35,105,106) and histone deacetylase (HDAC) (103,107,108), the epigenome of the 12h oscillator is marked by COMPASS-mediated histone methylation, specifically H3K4me3 at the proximal promoters. Loss of one of the essential subunits of COMPASS, RBBP5, abolished more than 90% of the hepatic ∼12h transcriptome, while having no effects on the core circadian clock. Knocking down RBBP5 in MEFs further dampened the cell-autonomous 12h oscillations, but not the circadian rhythms. We further expanded our findings to the epigenetic regulation of response to acute and chronic proteotoxic stress and found both increased RBBP5 promoter recruitment and H3K4me3 level are associated with the transcription activation of adaptive stress response genes. Subsequently, depletion of RBBP5 significantly dampened the adaptive transcriptional responses to various proteotoxic stresses.

In our study, we uncovered distinct roles for H3K4me3 and H3K9Ac/H3K27Ac in regulating the 12h oscillator and transcriptional responses to acute proteotoxic stress. While all three histone modifications—H3K4me3, H3K9Ac, and H3K27Ac—are traditionally linked to transcriptional activation (81,82,109), our findings suggest that H3K9Ac and H3K27Ac are dispensable for the transcriptional regulation of proteostasis under both basal (the 12h oscillator) and stress conditions. Interestingly, recent research by (110) demonstrated that among various epigenetic marks associated with transcriptional activation, the installation of H3K4me3 at promoters resulted in the most potent transcriptional activation. This not only underscores the causal role of H3K4me3 in gene activation but also highlights the differential impacts of specific histone modifications on transcription. We speculate that among the many histone modifications, H3K4me3 has been evolutionarily selected as the primary epigenetic modification for stress response due to its ability to trigger the most robust gene activation, thereby providing a rapid adaptive advantage under adverse environmental conditions. Future investigations are needed to further test this hypothesis.

The precise role of H3K4me3 in transcriptional regulation remains a topic of debate. Recent evidence suggests that H3K4me3 positively contributes to transcriptional elongation by regulating the pause-release of RNAP II, likely through the recruitment of the Integrator subunit INST11 (89). Our H3K4me3 proximity labeling not only supports this finding but also unexpectedly reveals increased recruitment of the SWI/SNF complex to H3K4me3 during the UPR in MEFs. Additionally, we confirmed the co-occurrence of RBBP5, H3K4me3, and the PBAF subunit BAF180 in chromatin binding in vivo in mouse liver. Interestingly, the temporal dynamics of SWI/SNF recruitment to H3K4me3 coincide with the recruitment of the Integrator complex but occur after the initial increase in RNAP II subunits, which are likely involved in transcription initiation immediately following stress. This suggests that contrary to the widely accepted view that SWI/SNF primarily facilitates transcription initiation by increasing chromatin accessibility and transcription factor binding to promoter, it may also play a crucial yet underexplored role in transcription elongation. Although the literature on this aspect is rather limited, some earlier studies hint at this possibility (111,112). Future research is needed to firmly establish a causal relationship between SWI/SNF function, 12h oscillator regulation, and the proteostasis stress response, as well as to uncover the underlying molecular mechanisms.

The physiological and pathological roles of mammalian COMPASS subunits, particularly RBBP5, remain incompletely understood (91). While extensive research has primarily focused on their contributions to organogenesis, early development, and neurodevelopmental disorders, our study significantly broadens the scope of RBBP5’s physiological roles by uncovering its involvement in proteostasis and hepatic metabolism. This expansion not only advances our understanding of RBBP5’s function but also opens new avenues for exploring its broader impact on cellular and systemic processes. Of particular interest is a recent study that identified heterozygous loss-of-function mutations in *RBBP5* in humans, which were primarily linked to neurodevelopmental disorders (113). This finding raises intriguing questions about the full spectrum of RBBP5’s physiological roles. Given our discovery of RBBP5’s involvement in metabolic regulation, it is tempting to hypothesize that individuals with RBBP5 loss-of-function mutations may also exhibit metabolic defects. Such a connection would have significant implications, suggesting that RBBP5 plays a multifaceted role in maintaining both neural and metabolic homeostasis. Future research should investigate whether these mutations influence metabolic pathways, potentially contributing to a broader spectrum of clinical symptoms in affected individuals. This line of inquiry could ultimately lead to a deeper understanding of how COMPASS subunits like RBBP5 integrate signals across different tissues to maintain overall physiological balance.

Finally, we propose that the 12h oscillator serves as an effective ’discovery tool’ for uncovering previously unknown mechanisms of proteostasis regulation. By leveraging this approach, we recently identified a novel XBP1s-SON axis, which links nuclear speckle LLPS dynamics with the UPR (26). Herein, using a similar approach, we identified RBBP5 as an epigenetic regulator of proteostasis. We believe that continued exploration of the 12h oscillator will unveil further hidden principles governing proteostasis.

## Acknowledgements

We would like to thank Dr. Sebastien Gingras (University of Pittsburgh, Department of Immunology) for the design of the targeting strategy and identification of the founders carrying the properly targeted Rbbp5-flox allele. We would like to thank Chunming Bi and Zhaohui Kou of the Mouse Embryo Services Core (University of Pittsburgh, Department of Immunology) for microinjection of zygotes and production of Rbbp5-flox founder mice. We would also like to thank Dr. Heather Ballance, Dr. Naveen GV Kumar, Saad Irfan, C. Dupont, Dr. Bugra Zengin and Casey Edwards for assistance with mice colony maintenance, dissection, intraperitoneal injection and genotyping. We would also like to thank Dr. Michiel Vermeulen for kindly sharing the pK19HispATurbo plasmid. For time course transcriptome study, the library generation was performed in the Health Sciences Sequencing Core at UPMC Children’s Hospital of Pittsburgh, Rangos Research Center. Services and instruments used in this project were graciously supported, in part, by the University of Pittsburgh, the Office of the Senior Vice Chancellor for Health Sciences, the Department of Pediatrics, the Institute for Precision Medicine, and the Richard K Mellon Foundation for Pediatric Research.

W.D. was supported by fellowship F31 AG080998 through NIA. B. Zhu was supported by grants 1DP2GM140924 and 1R21AG071893 through the NIH, and a grant from Richard King Mellon foundation. This research was supported in part by the University of Pittsburgh Center for Research Computing through the resources provided. Specifically, this work used the HTC cluster, which is supported by NIH award number S10OD028483. This research project was supported in part by the Pittsburgh Liver Research Centre supported by NIH/NIDDK Digestive Disease Research Core Center grant P30DK120531.

## Author contributions

Conceptualization, B.Z.; Methodology, M.J. and B.Z.; Investigation, S.K., M.S., W.D., A.C., H.L., H.W., A.P., I.S., S.L., Y.P., L.L., J.J.L., and B.Z.; Writing – Original Draft, B.Z.; Writing – Review & Editing, all authors; Funding Acquisition, S.L., and B.Z.; Resources, M.J., J.H.L and B.Z.; Supervision, M.J., J.H.L and B.Z.

## Data availability

All raw and processed sequencing data generated in this study have been submitted to the NCBI Gene Expression Omnibus (GEO; http://www.ncbi.nlm.nih.gov/geo/) under accession numbers: GSE276155, GSE276157 and GSE276159. All data needed to evaluate the conclusions in the paper are present in the paper and/or the Supplementary Materials.

