## Supplemental figures for "The SET1/COMPASS subunit RBBP5 orchestrates epigenetic control of global proteostasis and the 12h oscillator to safeguard metabolic and cellular homeostasis"

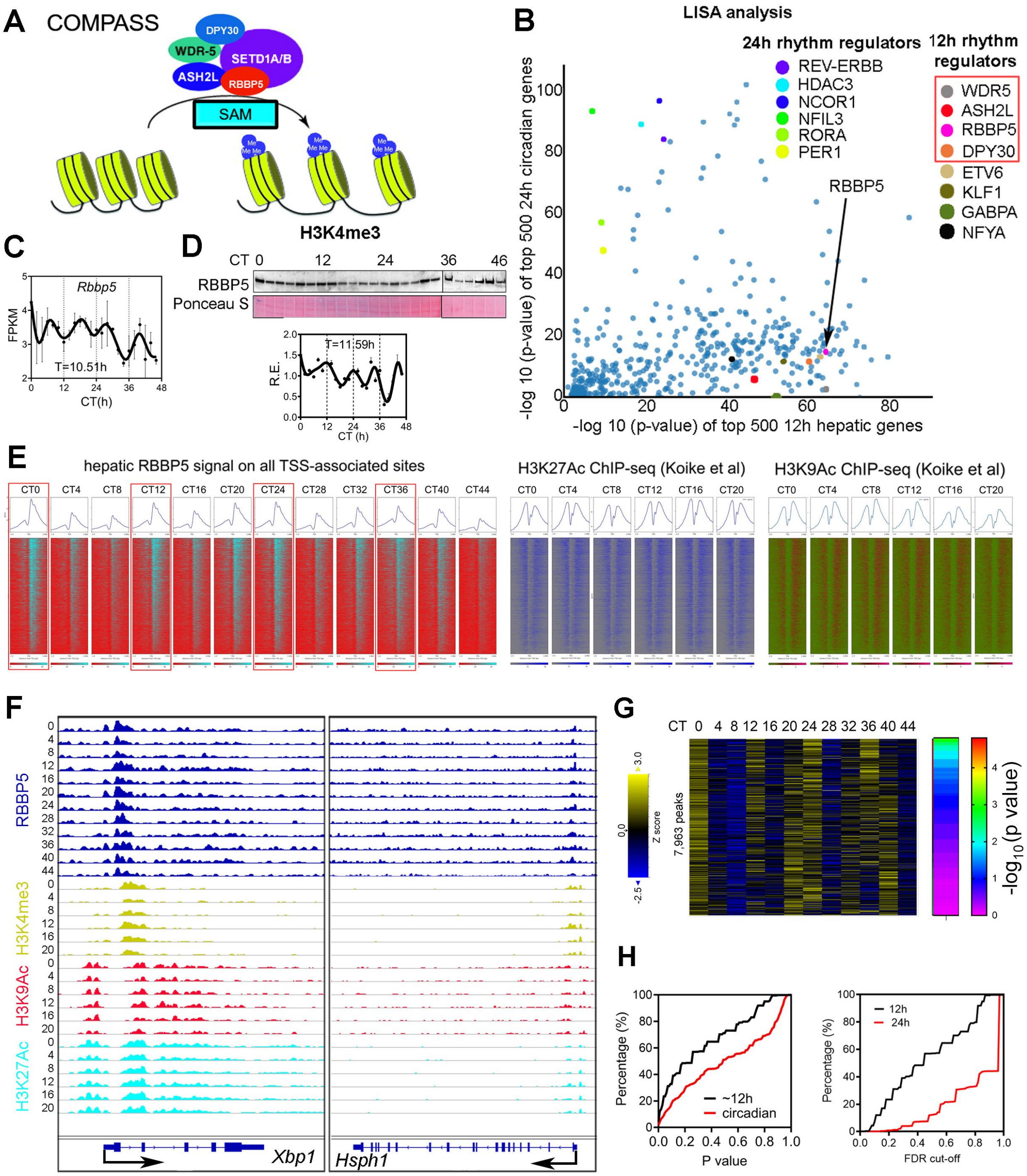


**Fig. S1. Global ~12h RBBP5 cistrome is associated with promoter-proximal ~12h H3K4me3 epigenome in mouse liver. (A)** COMPASS writes H3K4me3 using S-Adenosyl methionine (SAM) as the substrate. **(B)** LISA revealing putative TFs and co-regulators for hepatic top 500 most robust (the smallest p values by RAIN analysis) circadian (y-axis) and 12h (x-axis) genes. **(C)** Temporal hepatic *Rbbp5* expression assayed by RNA-seq. Period is calculated by the eigenvalue/pencil method. **(D)** Western blot and quantification (n=2) of temporal nuclear RBBP5 level in mouse liver at different CTs. Period is calculated by eigenvalue/pencil method. **(E)**

Heatmap showing RBBP5, H3K9Ac and H3K27Ac chromatin occupancy 1kb ± of TSS of 6,451 genes. **(F)** Snapshot of target genes selected for alignment of hepatic RBBP5, H3K4me3, H3K9Ac and H3K27Ac at different CTs. **(G)** Heatmap of temporal RBBP5 binding intensity for 7,963 binding sites, along with $-\log_{10}$ transformed P value for having 12h rhythm by RAIN. **(H)** Cumulative distribution of the number of ~12h and circadian RBBP5 binding sites ranked by p value or false discovery rate (FDR). Data: Mean ± S.E.M.

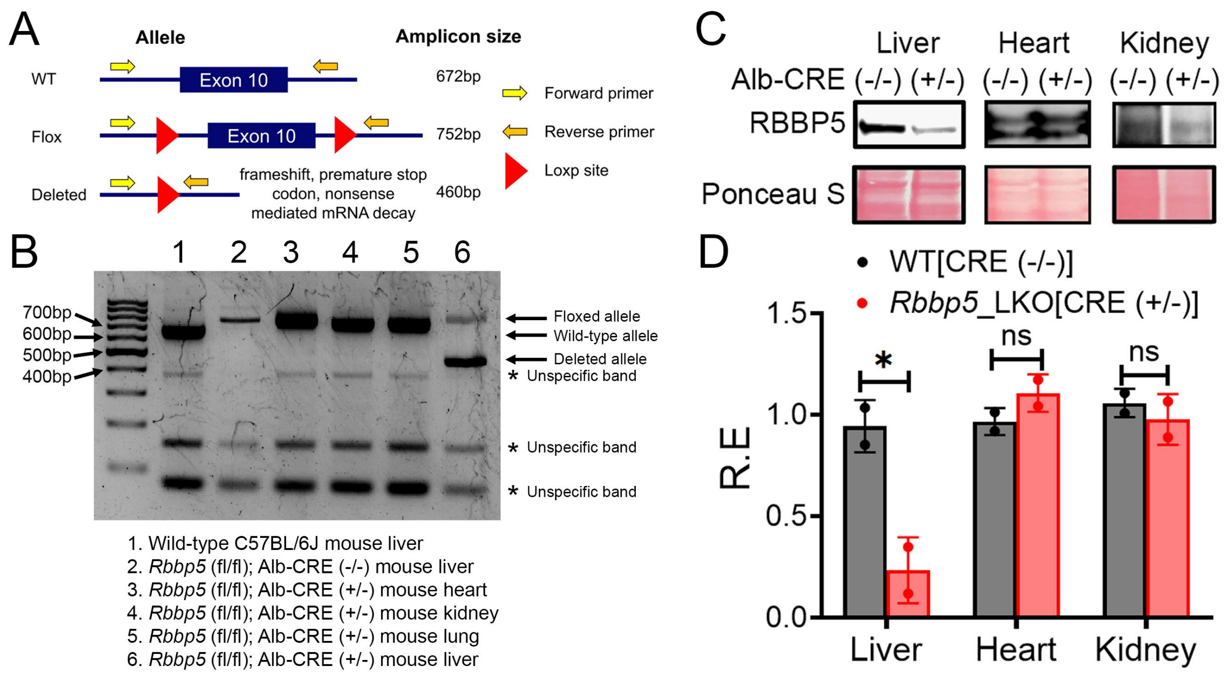


**Fig. S2. The generation of RBBP5 $^{LKO}$ mice. (A, B)** Expected **(A)** and actual genotyping result **(B)** for different mice. **(C, D)** Western blot **(C)** and quantification **(D)** of RBBP5 in different tissues in *Rbbp5* (fl/fl); Alb-CRE (+/-) and *Rbbp5* (fl/fl); Alb-CRE (-/-) mice. n=2 per genotype. Data: Mean ± S.E.M.

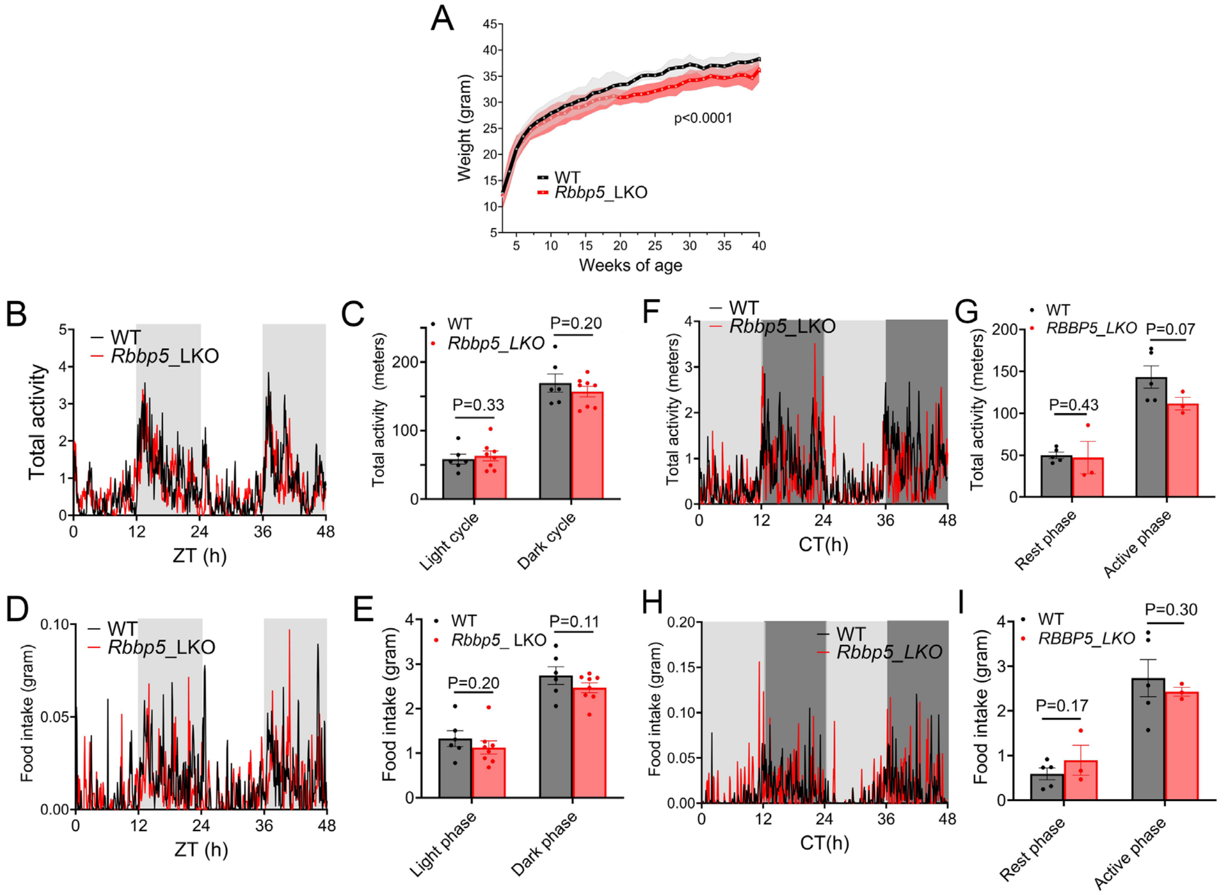


**Fig. S3. Liver-specific deletion of RBBP5 does not alter rhythmic locomotor activity nor fasting-feeding cycles in mice. (A)** Average body weight of male *Rbbp5* $^{Flox}$ and *Rbbp5* $^{LKO}$ mice at different ages. Mean ± 95% confidence interval. N=3~71 for *Rbbp5* $^{Flox}$ and n=6~54 for *Rbbp5* $^{LKO}$ mice at each week. P<0.0001 by One-way ANOVA. **(B)** Real-time locomotor activity monitoring in *Rbbp5* $^{Flox}$ and *Rbbp5* $^{LKO}$ mice under 12hr light/12hr dark conditions. **(C)** Averaged measurements within the light and dark phase as described in **B. (D)** Real-time measurement of food intake in *Rbbp5* $^{Flox}$ and *Rbbp5* $^{LKO}$ mice under 12hr light/12hr dark conditions. **(E)** Averaged measurements within the light and dark phase as described in **D.** n=6 for *Rbbp5* $^{Flox}$ and n=8 for *Rbbp5* $^{LKO}$ mice. **(F)** Real-time locomotor activity monitoring in *Rbbp5* $^{Flox}$ and *Rbbp5* $^{LKO}$ mice under constant dark conditions. **(G)** Averaged measurements within the rest and active phase as described in **F**. **(H)** Real-time measurement of food intake in *Rbbp5* $^{Flox}$ and *Rbbp5* $^{LKO}$ mice under constant dark conditions. **(I)** Averaged measurements within the rest and active phase as described in **H.** n=5 for *Rbbp5* $^{Flox}$ and n=3 for *Rbbp5* $^{LKO}$ mice. Data: Mean ± S.E.M.

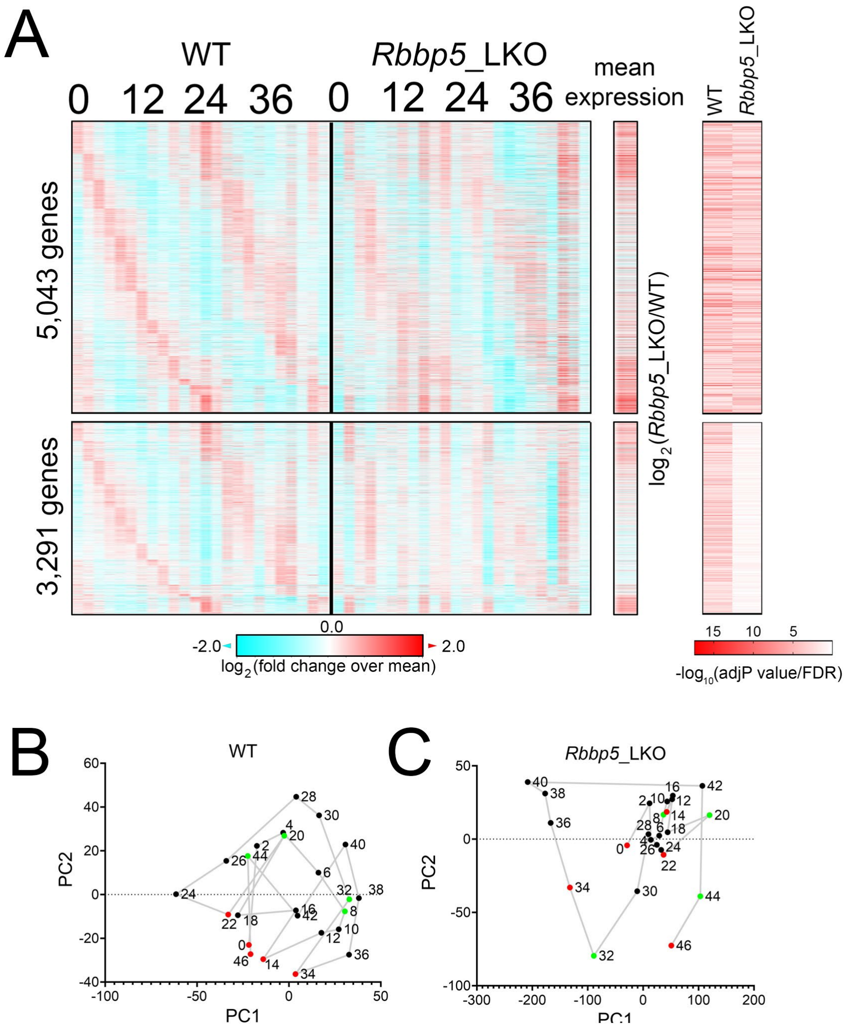


D

| Criterion | WT | *Rbbp5* _LKO | Shared |
|---|---|---|---|
| 10-12h (FDR<0.001) | 1,113 | 4 | 2 |
| 10-12h (FDR<0.01) | 3,549 | 61 | 13 |
| 10-12h (FDR<0.05) | 6,025 | 813 | 424 |
| 22-28h (FDR<0.01) | 6,324 | 5,456 | 2,939 |
| 22-28h (FDR<0.05) | 8,334 | 7,515 | 5,043 |

**Fig. S4. RBBP5 is an epigenetic regulator of the hepatic ~12h oscillator, but not the canonical ~24h circadian clock. (A)** Heat map of 8,340 circadian gene expression (FDR<0.05) in the liver of *Rbbp5* $^{Flox}$ and *Rbbp5* $^{LKO}$ mice, with 5,043 of them shared between the two, along with -$\log_{10}$ transformed adjP values for having 22-28h rhythm by RAIN. $\text{Log}_2$ normalized fold change of average gene expression across 48 hours between *Rbbp5* $^{Flox}$ and *Rbbp5* $^{LKO}$ mice was also shown. (**B, C**) PCA of hepatic temporal transcriptome in *Rbbp5* $^{Flox}$ (**B**) and *Rbbp5* $^{LKO}$ (**C**) mice. **(D)** A table listing the number of ~12h and ~24h circadian genes in *Rbbp5* $^{Flox}$ and *Rbbp5* $^{LKO}$ mice with different statistical thresholds.

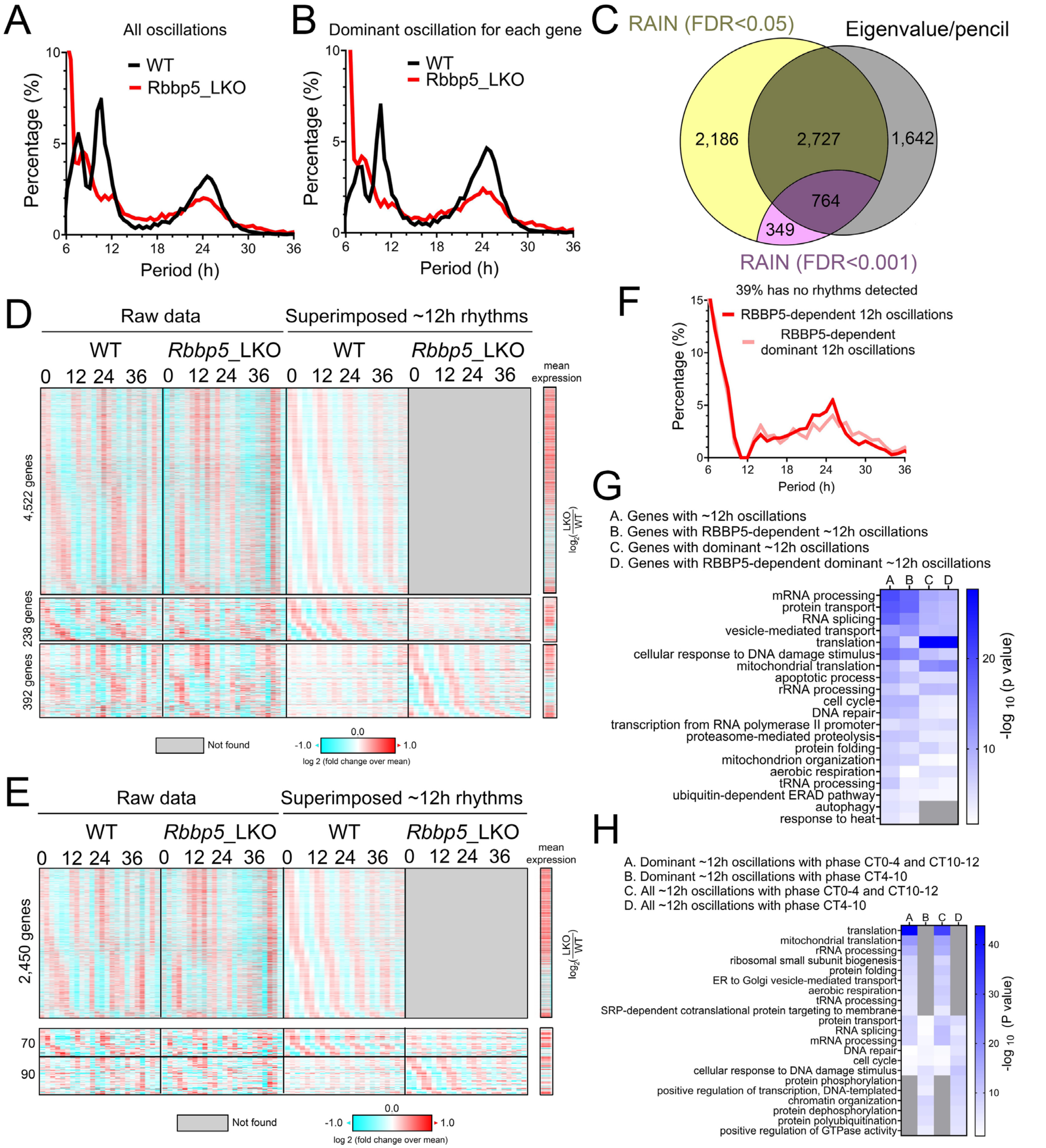


**Fig. S5. Eigenvalue/pencil method analysis of RBBP5-dependent hepatic ~12h transcriptome. (A, B)** Distributions of all **(A)** and dominant **(B)** oscillations uncovered by the eigenvalue/pencil method in *Rbbp5* $^{Flox}$ and *Rbbp5* $^{LKO}$ mice. **(C)** Venn diagram comparing unique and common ~12h transcriptomes uncovered by the eigenvalue/pencil and RAIN methods with two different FDR cut-offs. **(D)** Heat map of all ~12h gene expression (or lack thereof) in *Rbbp5* $^{Flox}$ and *Rbbp5* $^{LKO}$ mice with both raw data and superimposed ~12h rhythms shown. 4,522 ~12h genes were abolished, 238 ~12h genes dampened, and 392 ~12h genes were enhanced in *Rbbp5* $^{LKO}$ mice, respectively. $Log_2$ normalized fold change of average gene expression across

48 hours between *Rbbp5* $^{Flox}$ and *Rbbp5* $^{LKO}$ mice was also shown. **(E)** Heat map of dominant ~12h gene expression (or lack thereof) in *Rbbp5* $^{Flox}$ and *Rbbp5* $^{LKO}$ mice with both raw data and superimposed ~12h rhythms shown. 2,450 ~12h genes were abolished, 70 ~12h genes dampened, and 90 ~12h genes were enhanced in *Rbbp5* $^{LKO}$ mice, respectively. $Log_2$ normalized fold change of average gene expression across 48 hours between *Rbbp5* $^{Flox}$ and *Rbbp5* $^{LKO}$ mice was also shown. **(F)** The periods distribution in *Rbbp5* $^{LKO}$ mice of those ~12h genes originally identified in *Rbbp5* $^{Flox}$ mice but lost in *Rbbp5* $^{LKO}$ mice. **(G, H)** Heat map summary of GO analysis demonstrating the $-log_{10}$ transformed P values of different enriched pathways for all **(G)** and dominant **(H)** ~12h genes.

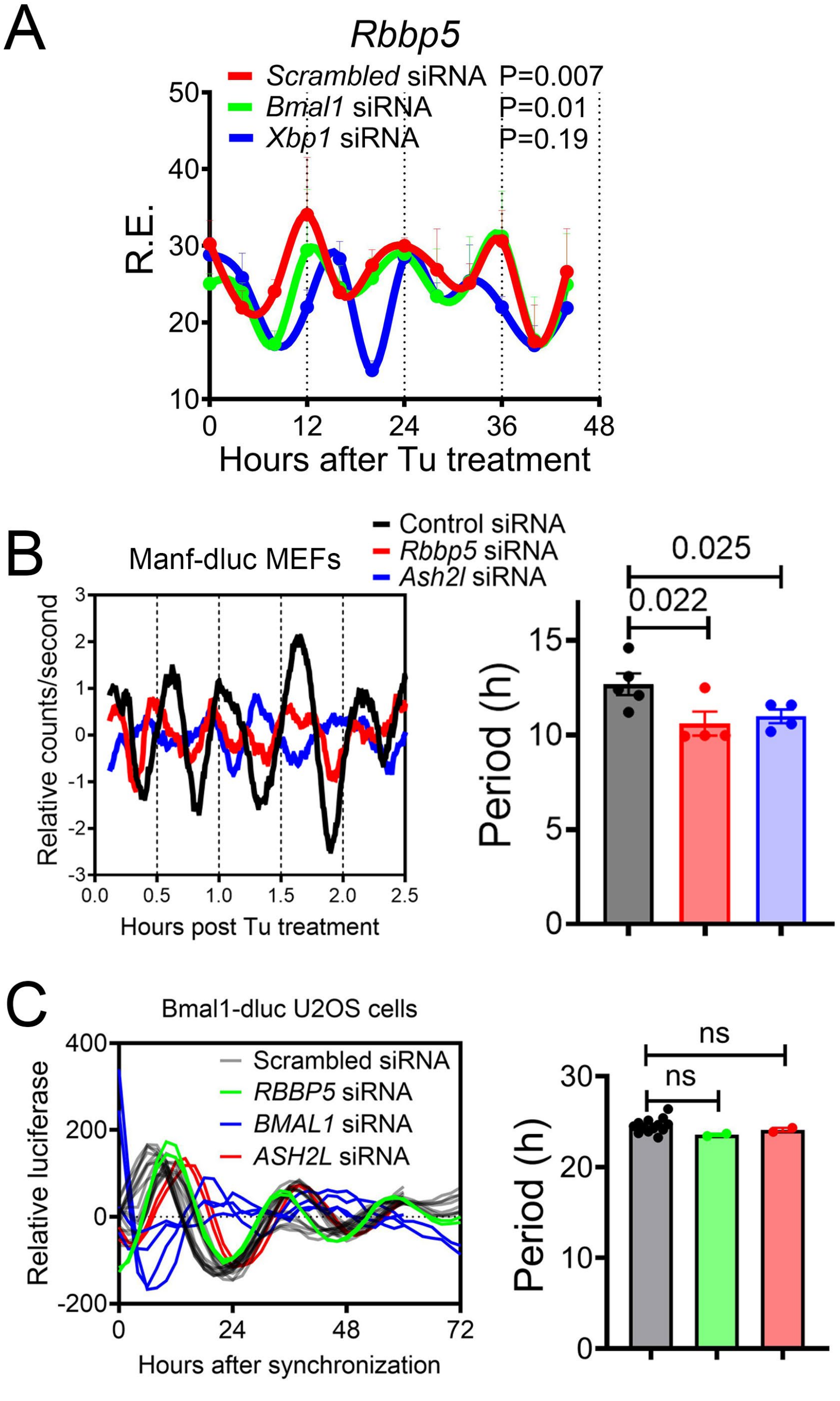


**Fig. S6. RBBP5 is a cell-autonomous epigenetic regulator of the ~12h oscillator, but not the canonical ~24h circadian clock. (A)** qPCR analysis of *Rbbp5* expression in Tu (25ng/ml)-synchronized MEFs with scrambled, *Bmal1* and *Xbp1* siRNAs. P values for having 12h rhythms were calculated by RAIN. **(B)** Real-time luminescence of MEFs expressing *Manf* promoter-driven dluc (23) transfected with different siRNAs and quantified periods. **(C)** Real-time luminescence traces of Bmal1-dluc U2OS cells transfected with different siRNAs as reported in (68). Data: Mean ± S.E.M.

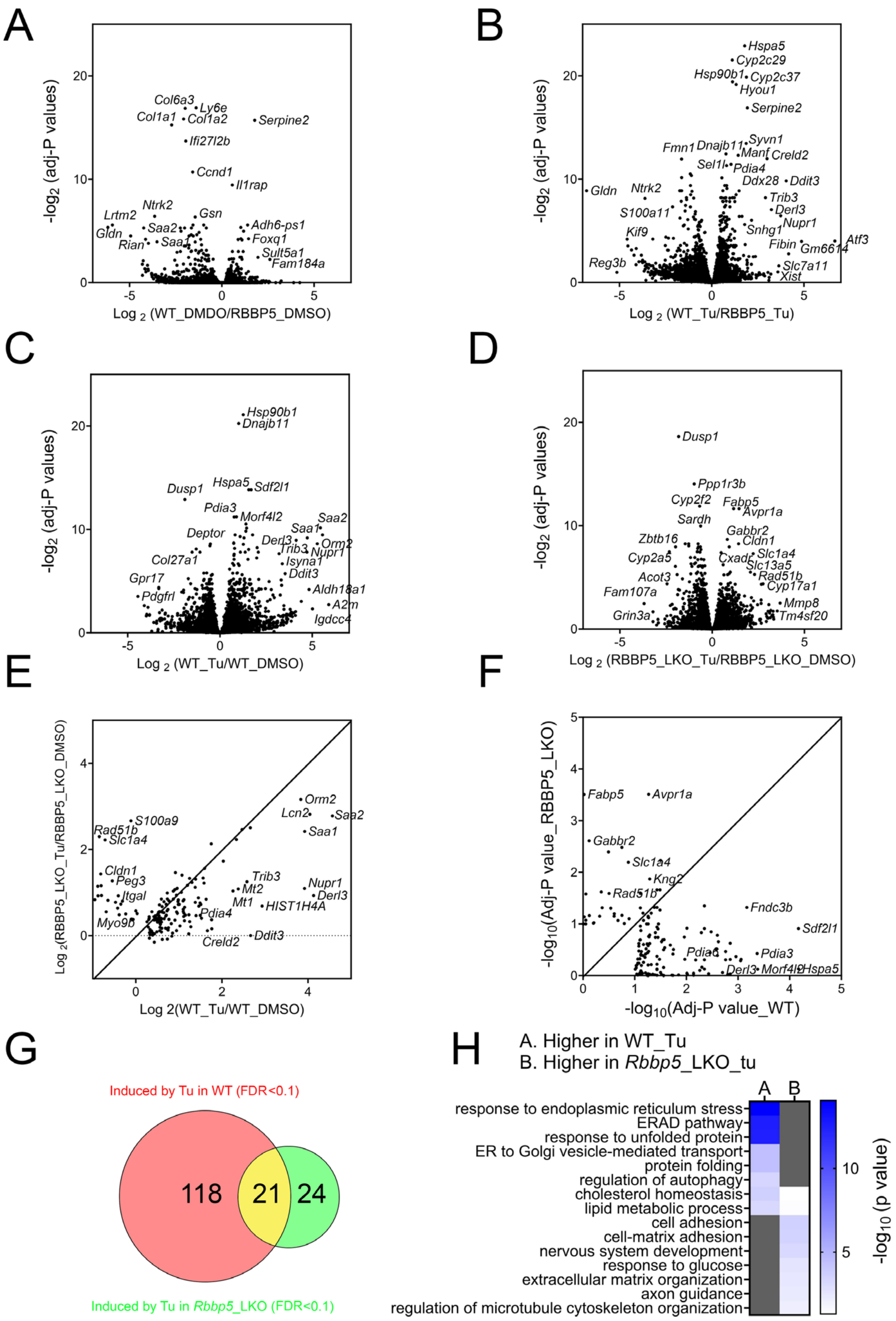


**Fig. S7. RBBP5 regulates the hepatic transcriptional response to proteotoxic stress. (A-D)** Volcano plot illustrating the log 2 normalized fold change vs -log2 transformed adjusted p values for different comparisons. **(E)** Scatter plot comparing the log 2 transformed fold change of gene expression by Tu in *Rbbp5* $^{Flox}$ (x- axis) and *Rbbp5* $^{LKO}$ (y-axis) mice. **(F)** Scatter plot comparing the -log 10 transformed adjusted p values for gene expression induced by Tu in *Rbbp5* $^{Flox}$ (x- axis) and *Rbbp5* $^{LKO}$ (y-axis) mice. **(G)** Venn diagram comparing distinct and common genes induced by Tu in *Rbbp5* $^{Flox}$ and *Rbbp5* $^{LKO}$ mice with FDR<0.1. **(H)** Heat map summary of GO analysis demonstrating the -log$_{10}$ transformed P values of different enriched pathways for different groups of genes.

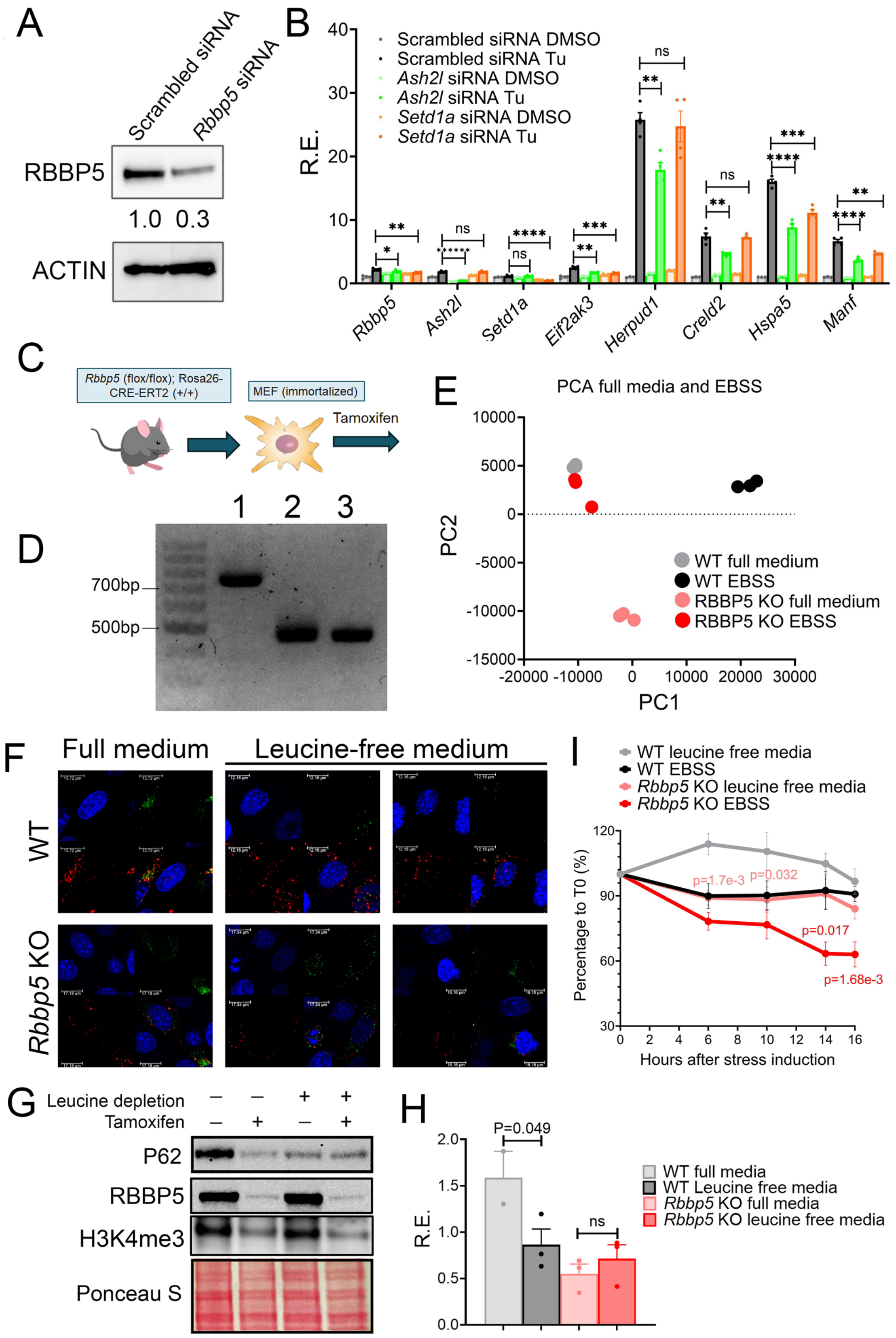
A
Scrambled siRNA
Rbbp5 siRNA
RBBP5
1.0 0.3
ACTIN
B
Scrambled siRNA DMSO
Scrambled siRNA Tu
Ash2l siRNA DMSO
Ash2l siRNA Tu
Setd1a siRNA DMSO
Setd1a siRNA Tu
R.E.
Rbbp5
Ash2l
Setd1a
Eif2ak3
Herpud1
Creld2
Hspa5
Manf
C
Rbbp5 (flox/flox); Rosa26-CRE-ERT2 (+/+)
MEF (immortalized)
Tamoxifen
D
1 2 3
700bp
500bp
E
PCA full media and EBSS
PC1
PC2
WT full medium
WT EBSS
RBBP5 KO full medium
RBBP5 KO EBSS
F
Full medium
Leucine-free medium
WT
Rbbp5 KO
I
WT leucine free media
WT EBSS
Rbbp5 KO leucine free media
Rbbp5 KO EBSS
Percentage to T0 (%)
Hours after stress induction
p=1.7e-3
p=0.032
p=0.017
p=1.68e-3
G
Leucine depletion
Tamoxifen
P62
RBBP5
H3K4me3
Ponceau S
H
P=0.049
ns
R.E.
WT full media
WT Leucine free media
Rbbp5 KO full media
Rbbp5 KO leucine free media

1568

**Fig. S8. RBBP5 is required for transcriptional responses to diverse proteotoxic stresses. (A)** Western blot of RBBP5 in MEFs transfected with *scrambled* or *Rbbp5* siRNA for 48 hours. **(B)** qPCR of different genes in MEFs transfected with scrambled, *Ash2l* or *Setd1a* siRNAs and treated with DMSO or Tu (100ng/ml) for 8 hours. **(C, D)** Isolation and immortalization of *Rbbp5* (fl/fl) ROSA26-CreERT2 (+/+) MEFs **(C)**, which were treated with vehicle (WT) or tamoxifen (*Rbbp5* KO) for 5 or 7 days and genotyping **(D)** to confirm the deletion of floxed *Rbbp5* allele. (**E**) PCA plot of transcriptome of ROSA26-CreERT2 (+/+) MEFs treated with vehicle (WT) or tamoxifen (*Rbbp5* KO) followed by treatment with full DMEM media or EBSS for 16 hours. **(F)** Representative images of autophagic flux in mCherry-GFP-LC3-expressing *Rbbp5* (fl/fl) ROSA26-CreERT2 (+/+) MEFs treated with vehicle (WT) or tamoxifen (*Rbbp5* KO) followed by treatment with full DMEM media or leucine-free media for 16 hours. **(G, H)** Western blot of different proteins **(G)** and quantification **(H)** of p62 in *Rbbp5* (fl/fl) ROSA26-CreERT2 (+/+) MEFs treated with vehicle (WT) or tamoxifen (*Rbbp5* KO) followed by treatment with full DMEM media or leucine free media for 16 hours**. (I)** Percentage of cells normalized to before treatment for mCherry-GFP-LC3-expressing *Rbbp5* (fl/fl) ROSA26-CreERT2 (+/+) MEFs treated with vehicle (WT) or tamoxifen (*Rbbp5* KO) followed by treatment with full DMEM media, leucine-free media or EBSS for up to 16 hours. Data: Mean ± S.E.M.

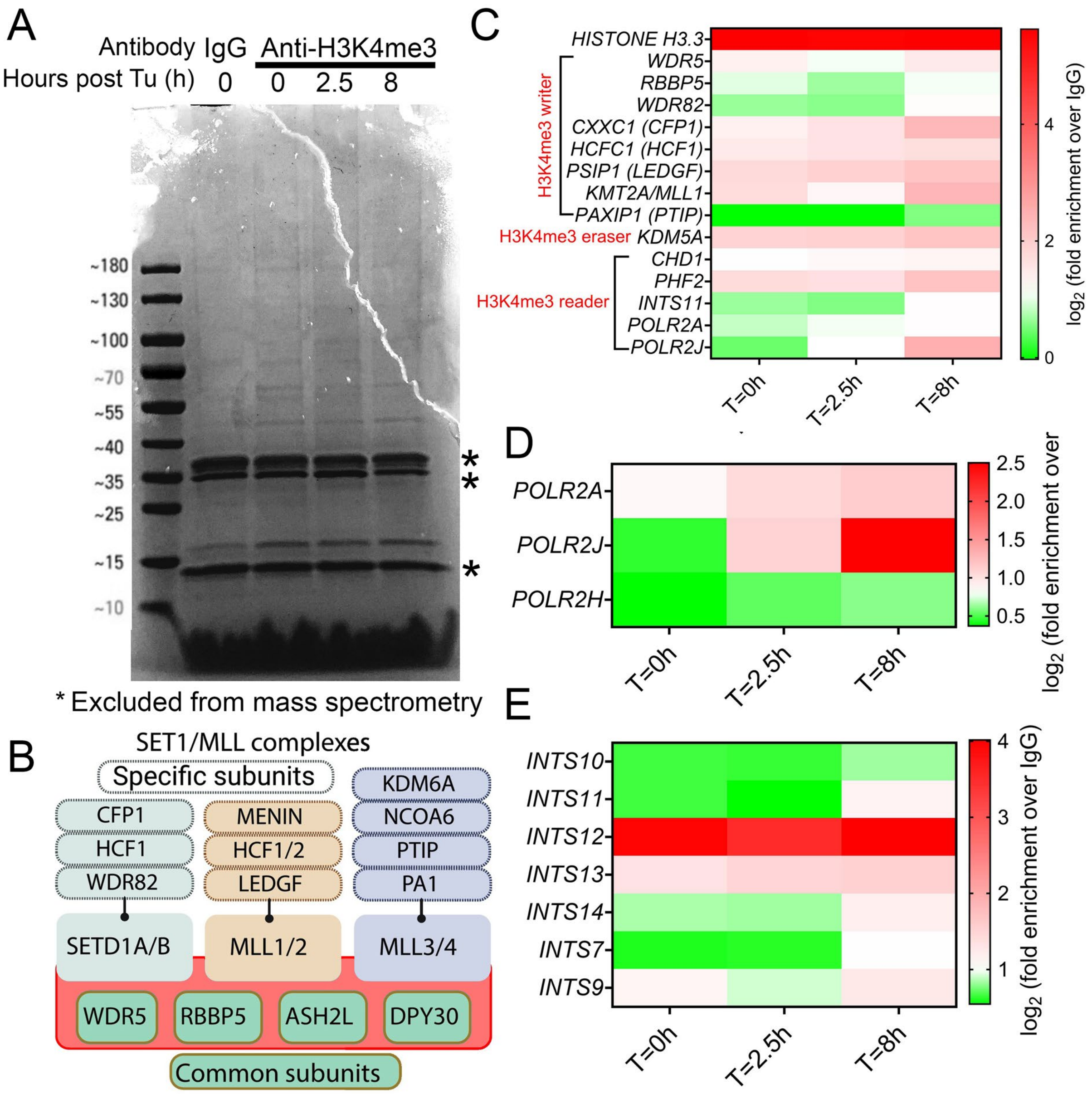


**Fig. S9. Proximity labeling reveals H3K4me-associated proteomic architecture of the proteostatic stress response.** (**A**) Imperial blue staining of total biotinylated proteins in MEF treated with 100ng/ml Tu for 0, 2.5 or 8h before subject to anti-IgG and anti-H3K4me3 antibody-based proximity labeling. Unspecific bands (in a different gel) denoted by asterisks are excluded from mass spectrometry analysis by gel removal. (**B**) A diagram illustrating common and specific subunits of mammalian COMPASS complex. (**C-E**) Heatmap of fold enrichment of different subunits in COMPASS (**C**), RNAP II (**D**) or Integrator Complex € (over IgG control) at different times after ER stress.

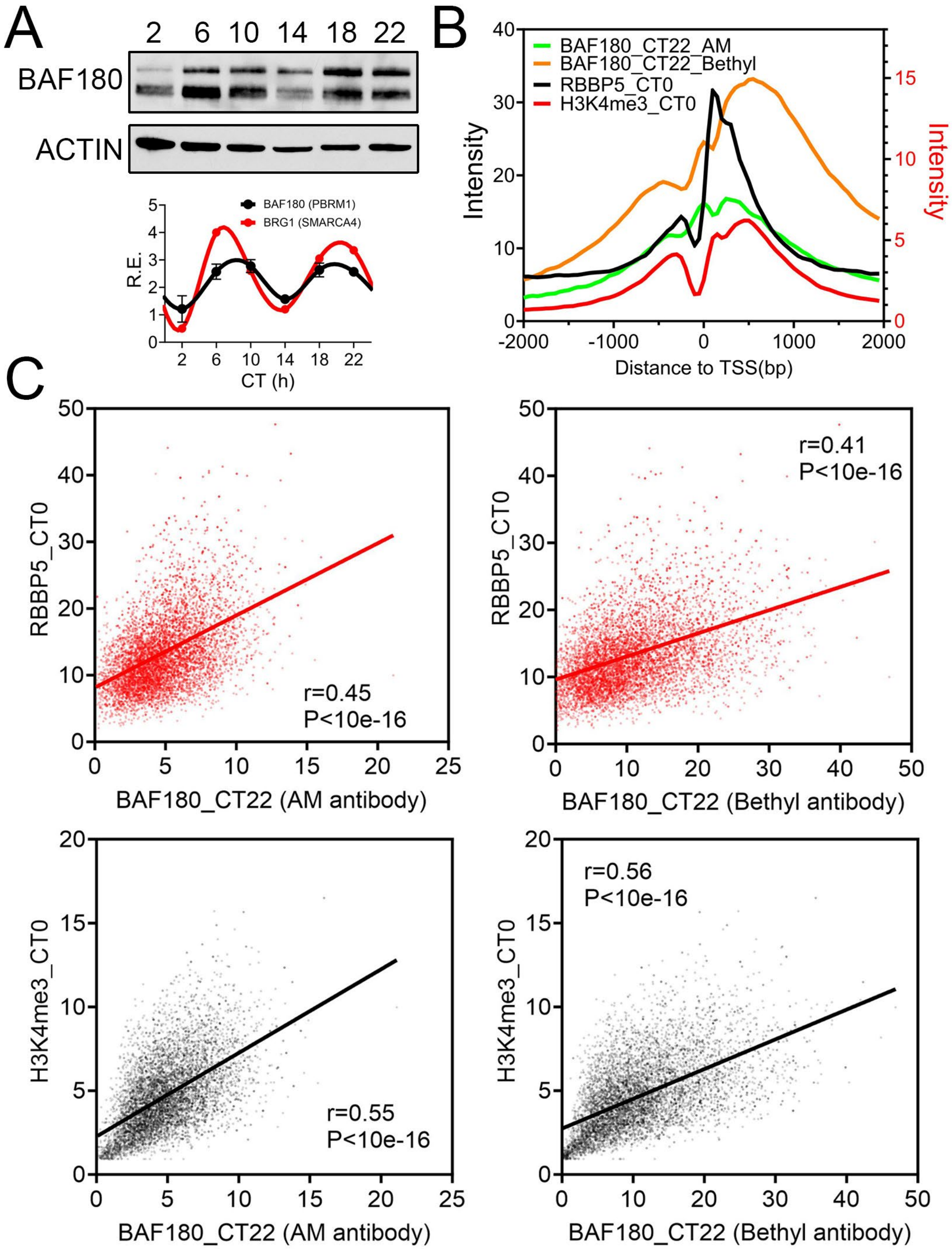


**Fig. S10. BAF180 cistrome colocalizes with those of RBBP5 and H3K4me3 in mouse liver.** (**A**) Western blot and quantification of nuclear BAF180 in the liver of wild-type mouse liver at different CTs. The quantification of BRG1 is based upon the original blot reported in Fig. 5E in (36). (**B**) Quantification of RBBP5, H3K4me3 and BAF180 chromatin occupancy 2kb ± of TSS of 6,451 genes. (**C**) Scatter plot comparing BAF180 binding signal with those of RBBP5 and H3K4me3 2kb downstream of TSS of 6,451 genes.

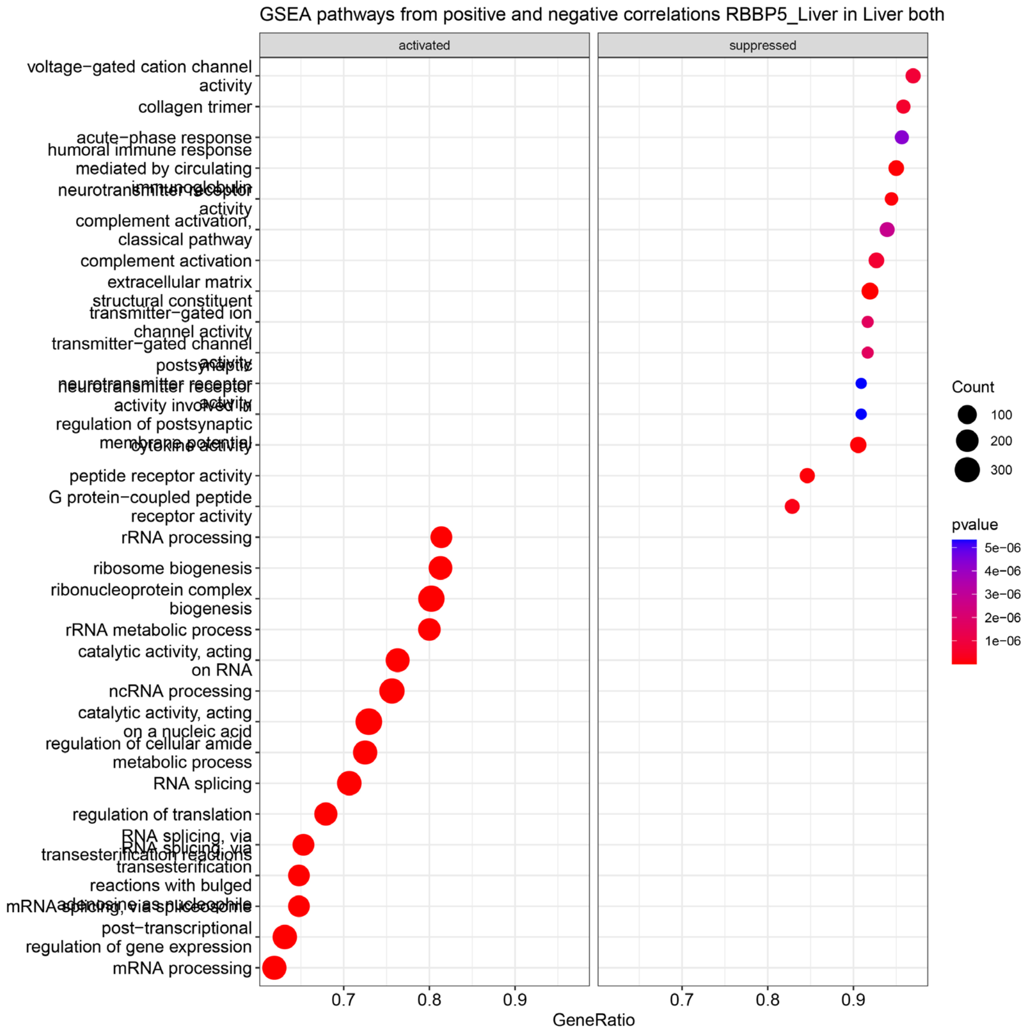


**Fig. S11. RBBP5 expression is positively correlated with those involved in proteostasis and mRNA metabolism and negatively associated with those implicated in immune response in human liver.** Data are from the human GD-CAT dataset (98).

### Supplemental Tables

**Table S1. FPKM quantification of temporal hepatic RBBP5 ChIP-seq in wild-type mouse liver.**

**Table S2. FPKM quantification of temporal hepatic RNA-seq in *Rbbp5* $^{Flox}$ and *Rbbp5* $^{LKO}$ mice.**

**Table S3. RAIN analysis of 10~12h hepatic transcriptome identified in *Rbbp5* $^{Flox}$ and *Rbbp5* $^{LKO}$ mice.**

Tab 1: 10~12h oscillations identified in *Rbbp5* $^{Flox}$ mice.

Tab 2: 10~12h oscillations identified in *Rbbp5* $^{LKO}$ mice.

**Table S4. Eigenvalue/pencil analysis of oscillating hepatic transcriptome identified in *Rbbp5* $^{Flox}$ and *Rbbp5* $^{LKO}$ mice.**

Tab 1: All oscillations identified in *Rbbp5* $^{Flox}$ mice.

Tab 2: All dominant oscillations identified in *Rbbp5* $^{Flox}$ mice.

Tab 3: All 10~13h oscillations identified in *Rbbp5* $^{Flox}$ mice.

Tab 4: All dominant 10~13h oscillations identified in *Rbbp5* $^{Flox}$ mice.

Tab 5: All oscillations identified in *Rbbp5* $^{LKO}$ mice.

Tab 6: All dominant oscillations identified in *Rbbp5* $^{LKO}$ mice.

Tab 7: All 10~13h oscillations identified in *Rbbp5* $^{LKO}$ mice.

Tab 8: All dominant 10~13h oscillations identified in *Rbbp5* $^{LKO}$ mice.

**Table S5. RNA-seq quantification of hepatic transcriptome in *Rbbp5* $^{Flox}$ and *Rbbp5* $^{LKO}$ mice with or without Tu injection.**

Tab 1: Raw counts

Tab 2: CPM quantification

Tab 3: FKPM quantification

**Table S6. Differentially expressed gene (DEG) analysis of hepatic transcriptome in *Rbbp5* $^{Flox}$ and *Rbbp5* $^{LKO}$ mice with or without Tu injection.**

**Table S7. RNA-seq quantification (TPM) of transcriptome in WT and *Rbbp5* KO MEFs in response to EBSS treatment**

**Table S8. Differentially expressed gene (DEG) analysis of WT and *Rbbp5* KO MEFs in response to EBSS treatment**

**Table S9. Log 2 normalized counts of 239 proteins identified by H3K4me3 proximity labeling in MEFs that showed the highest enrichment at T=8h**
